# Sensory neurons gate a cortical plasticity window for immune encoding

**DOI:** 10.64898/2026.11.02.763631

**Authors:** Zachary H. Gursky, Shi Tang, Min Wu, Chenxin Liu, Shao-Yu Fang, Emory K. Farrell, Suganya Subramanian, Amanda Wu, Patricia A. Colom Díaz, Amira Ibrahim, Stefano Berto, Alicia Che, Emilia Favuzzi

**Author notes:** These authors contributed equally to this work (Zachary H. Gursky, Shi Tang).

## Abstract

Activity-dependent plasticity allows brain circuits to encode experience, but this has been established with brief stimuli. Immune responses develop over days and are represented in the insular cortex (IC). How the cortex encodes experiences with this temporal structure is unknown. Here we show that Trpv1+ sensory neurons open a multi-day cortical plasticity window matched to the immune response. Following peripheral immune challenges, Parvalbumin-expressing inhibitory interneurons are activated first, restricting plasticity. Vasoactive intestinal peptide-expressing interneurons respond two days later, disinhibiting IC excitatory neurons. Weaker challenges trigger the first phase but not the second, and preventing the transition blocks immune encoding, whereas lowering inhibition beforehand does not. These results identify a body-to-brain mechanism that gates cortical plasticity on the timescale of slowly evolving physiological experience.

## Main Text

Memories of the external world and of the body’s internal states shape adaptations to biological challenges. The brain encodes memories through engrams, operationally defined as sparse neuronal ensembles recruited during an experience whose reactivation is sufficient to elicit recall. Although engrams were first established for exteroceptive experiences, such as aversive memories (*1*, *2*), they have more recently been shown to extend to the physiological domain. Reactivating thermosensory hippocampal engrams reinstates whole-body thermoregulatory responses (*3*), and reactivating immune engrams formed in the insular cortex (IC) during peripheral immune challenges reinstates organ-specific inflammatory states weeks after recovery (*4*). Unlike the discrete exteroceptive stimuli that drive classical engram formation, peripheral immune challenges unfold over days, engaging the nervous system through sustained afferent transmission of peripheral immune information (*5–8*). The encoding of immune experiences as engrams implies circuit mechanisms capable of bridging this timescale: peripheral signals must reach the IC throughout the challenge, and cortical circuits must be configured to encode them across this extended window. Neither the visceral afferent pathways that convey these signals to the IC nor the intracortical mechanisms that permit their encoding are known. Identifying these mechanisms would reveal how the brain meets the requirements for constructing representations of slowly evolving bodily states.

The IC is a cortical hub for interoceptive processing. It integrates physiological signals into representations that guide autonomic and behavioral responses (*9*). Whether a circuit is plastic and can encode new information, including the recruitment of neurons into engrams, depends on the state of its local inhibitory network. Within the IC and throughout the cortex, inhibitory interneurons modulate pyramidal cell excitability, enforce engram sparsity, and dynamically constrain plasticity windows (*10–14*). Parvalbumin-expressing (PV+) interneurons provide the dominant source of perisomatic inhibition and exert powerful control over pyramidal cell firing, while somatostatin-positive (SST+) interneurons perform slower, context-dependent dendritic inhibition (*13*, *15*). Both cell types are themselves plastic, adapting their synaptic connectivity and activity states to behavioral context (*16–22*). PV+ interneurons in particular can occupy distinct activity states, transitioning between high- and low-activity configurations that differentially govern the network’s capacity for experience-dependent plasticity (*16–18*). This transition is controlled by vasoactive intestinal peptide (VIP)-expressing interneurons, which specialize in disinhibitory control and suppress SST+ and PV+ interneurons (*16*, *23*). The activity state of PV+ interneurons is further regulated by perineuronal nets (PNNs), extracellular matrix structures that condense around PV+ cell bodies. PNN assembly stabilizes PV+ interneurons in a high-activity, low-plasticity configuration, while their dissolution reopens windows of enhanced plasticity (*16*, *17*, *20*, *24*, *25*). Together, these mechanisms determine when cortical circuits are available to encode new experience.

Here we identify a body-to-brain pathway for cortical immune encoding. We find that Trpv1+ sensory neurons first lead to the activation of IC PV+ interneurons, restricting plasticity, and then trigger a gradual circuit reconfiguration that culminates in excitatory neuron disinhibition. We demonstrate that the reconfiguration to a disinhibited state is governed by VIP+ interneurons and is required for immune engram formation. These findings establish multi-day inhibitory circuit plasticity as the mechanism by which the IC matches the slow timescale of peripheral immune experiences.

## Gut inflammation reorganizes brain-wide connectivity and recruits IC PV+ interneurons

To follow the brain’s response across the days over which a peripheral immune challenge develops, we used dextran sulfate sodium (DSS)-induced colitis, a model in which immune encoding has been demonstrated (*4*, *26*). Mice given 5% DSS developed the hallmarks of acute gut inflammation within four days: soft feces with occasional blood, colon shortening, CD45+ immune cell recruitment to the colon mucosa, and histopathological features of inflammation (Fig. 1A and fig. S1). To identify the brain regions recruited during the progression of gut inflammation, we used functional ultrasound imaging (fUSI). We assayed functional connectivity between anatomically defined regions of interest in two coronal imaging volumes (anterior and posterior scans that together spanned the mid-posterior IC), in control and DSS mice at baseline, day 2, and day 4 of DSS treatment (Fig. 1B). Functional connectivity remained stable in control mice throughout the experiment but reorganized in DSS mice as colitis progressed (fig. S2A). DSS mice differed little from baseline at day 2 but changed markedly by day 4 (Fig. 1, C and D; fig. S2, A to E). This reorganization did not spread evenly across the brain. Instead, a subset of regions gained connectivity above the network average. When we ranked every region by the increase in its overall connectivity, the IC was the most recruited region in the anterior plane (Fig. 1E and fig. S2F; agranular insula, AI, Allen brain nomenclature). In the posterior plane, the IC sat at the top of a cluster of strongly recruited limbic and temporal regions that included the hippocampal formation, entorhinal and perirhinal cortex, and the amygdala (Fig. 1F and table S1). The IC strengthened its coupling with distinct partners in each plane: with somatomotor, somatosensory, anterior cingulate, striatal, and pallidal regions in the anterior scan, and with hippocampal, retrohippocampal, perirhinal, and amygdalar regions in the posterior scan, together with hypothalamic regions in both (fig. S2D). These results show that gut inflammation progressively reorganized functional coupling in a set of brain circuits, with the IC emerging as a hub of this reorganization.

**Fig. 1.**
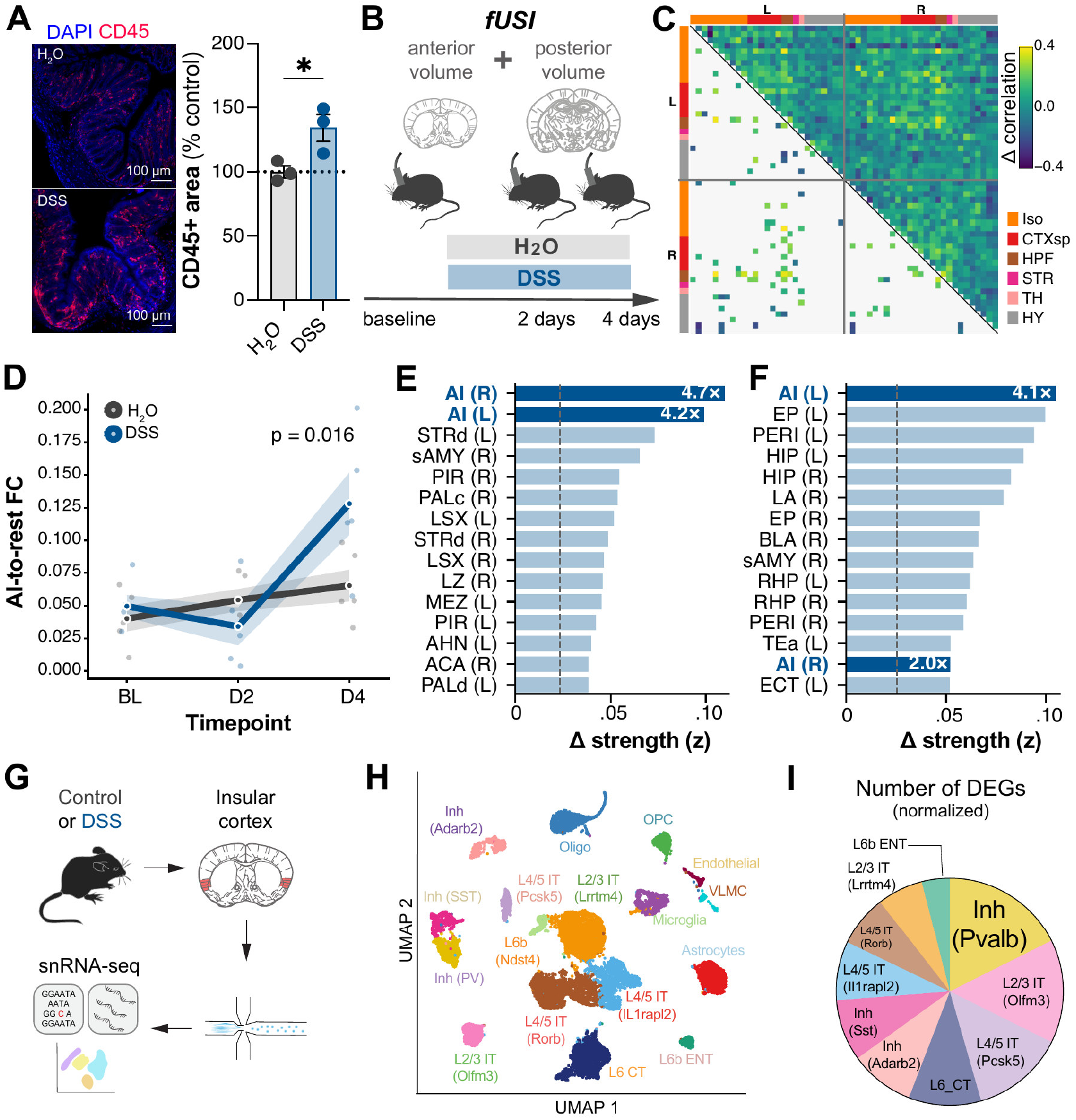
The IC and its PV+ interneurons are the regions and neuronal type most responsive to gut inflammation. **(A)** Representative images and quantification of CD45+ cell infiltration in the colon mucosa in H_2_O and 4-day DSS mice (n = 3 mice/group). Data in A are mean ± SEM, *p<0.05; unpaired t-test. **(B)** Schematic of the fUSI experiment (n = 5 mice/group). **(C)** Change in pairwise functional connectivity (Δ group-mean Pearson r), Day 4 minus baseline, DSS mice, posterior plane. Upper triangle, all pairs; lower triangle, pairs changed in DSS (paired t-test on Fisher z, p < 0.05). Ordered by hemisphere (L, R), then Allen macroregion (abbreviations in fig. S2). **(D)** Core insula (AI) connectivity with all other ROIs, posterior plane, at baseline (BL), Day 2 (D2) and Day 4 (D4). Lines: means; bands: ± SEM; dots: individual mice. Baseline, H_2_O versus DSS: unpaired t-test, p = 0.49; H_2_O across days: repeated-measures ANOVA, p = 0.16; DSS, D4 versus BL: paired t-test, p = 0.016. **(E-F)** Top 15 regions by increase in nodal strength (mean Fisher z correlation with all other regions), BL to D4, DSS mice, anterior (E; of 44) and posterior (F; of 54) planes. Dashed lines: network average; AI: dark blue, fold change above network average is indicated. **(G)** Schematic of snRNA-seq experiment. **(H)** UMAP of snRNA-seq dataset, annotated by IC cell type. **(I)** Number of differentially expressed genes (DEGs) per neuronal cell type, normalized to the number of nuclei recovered for that type.

Since fUSI established progressive macroscale IC recruitment, we next asked which cellular events lead to this recruitment. We performed single-nucleus RNA sequencing (snRNA-seq) of IC cells from control and DSS-treated mice at day 2 (Fig. 1, G and H, and fig. S3), preceding the connectivity peak. Among all IC neuronal types, PV+ inhibitory neurons exhibited the largest number of normalized differentially expressed genes (DEGs) in response to DSS (Fig. 1I). Cell-cell communication analysis revealed an expansion of outgoing signaling from PV+ interneurons in DSS conditions (fig. S3, D to F), indicating that PV+ cells in the IC not only respond most strongly at the transcriptional level but also reorganize their intercellular communication following immune challenge.

## Gut inflammation reconfigures IC inhibitory circuits into a plasticity-permissive state

Having identified PV+ interneurons as the most responsive population, we asked how their activity changes over the course of colitis. Because PV+ and SST+ interneurons bidirectionally gate engram size in classical memory systems (*10–12*, *14*), we tracked the activation kinetics of both using endogenous c-Fos as a proxy for neuronal activation. At 24 hours after DSS onset, neither PV+ nor SST+ cell populations showed changes in the fraction of active neurons (fig. S4A). By day 2 of DSS, the proportion of active PV+ cells had significantly increased compared to water-drinking controls (Fig. 2, A and B), while SST+ interneurons showed no significant changes (fig. S4B), indicating a selective early activation of PV+ interneurons. By day 4, the time point of most robust IC functional connectivity change (Fig. 1, C to F), the fraction of active PV+ cells fell significantly below baseline (Fig. 2, A and C). PV protein expression, an established activity-dependent molecular readout that inversely correlates with network plasticity (*16*, *17*), tracked the PV+ c-Fos+ fraction across both time points (Fig. 2D). The fraction of active SST+ interneurons was also reduced at the 4-day time point (fig. S4C). The proportion of active PV+ interneurons was unchanged in the primary visual cortex (V1, fig. S4D), ruling out a non-specific systemic inflammatory effect on cortical inhibitory networks. Inhibitory responses also scaled with disease severity, as DSS doses that produced milder colitis failed to recapitulate the biphasic PV+ cell dynamics (fig. S1C and fig. S4, E and F). Similarly, the attenuated colitis severity previously reported in female mice (*27*) triggered PV+ activation but not the 4-day inactivation (fig. S4, G to I), prompting the use of male mice in subsequent DSS experiments.

**Fig. 2.**
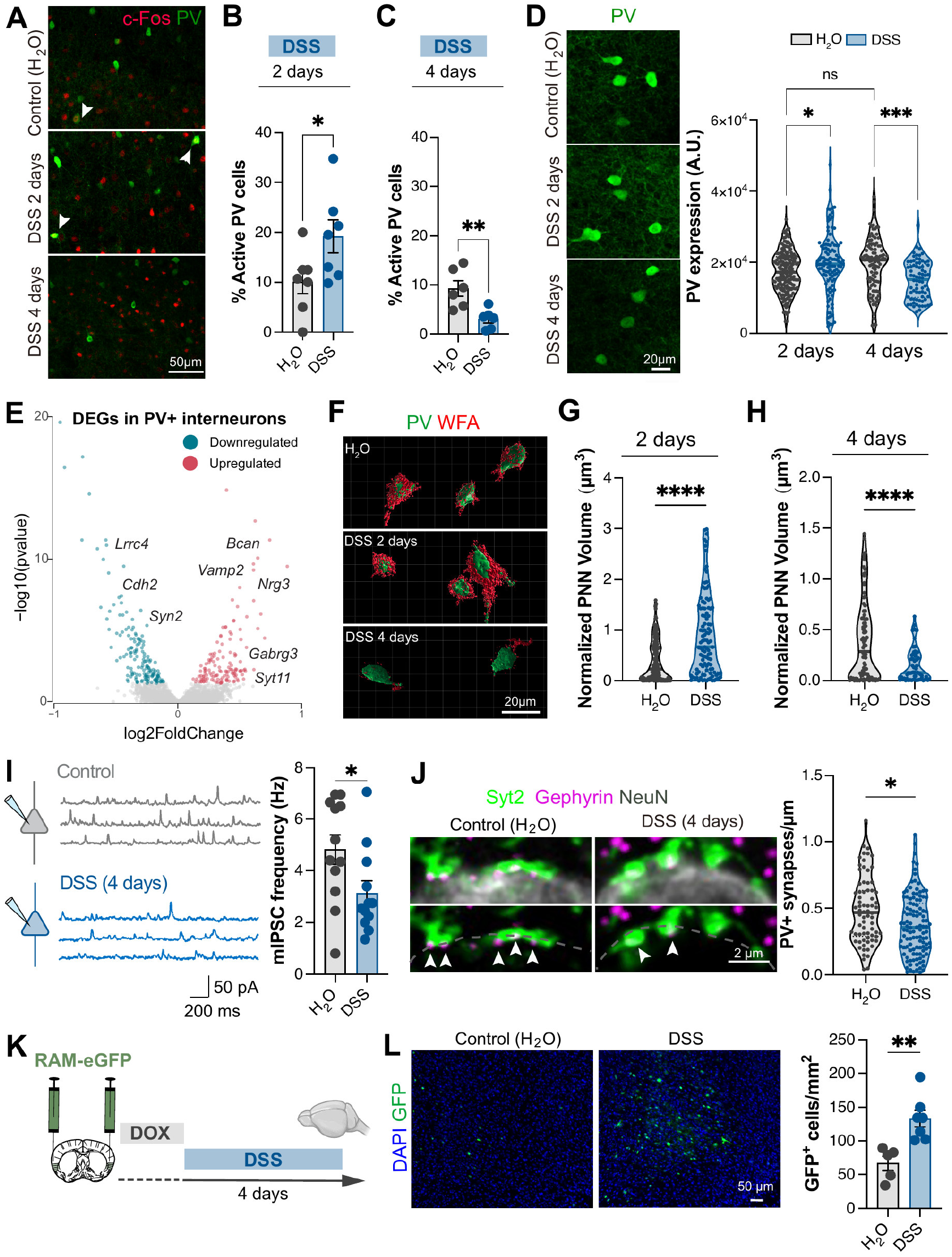
A biphasic reconfiguration of IC inhibitory circuits precedes IC plasticity and recruitment during gut inflammation. (A-C) Representative images (A) and fraction (B-C) of c-Fos+PV+ interneurons at 2 (B) and 4 (C) days of DSS (n = 6-7 mice/group). Arrowheads: colocalization. **(D)** PV intensity at 2 and 4 days of DSS (n = 74-188 cells from 5-7 mice/group). **(E)** DEGs in PV+ interneurons at 2 days of DSS. **(F-H)** 3D reconstructions (F) and quantification (G-H) of PNN volume (WFA) around PV+ interneurons at 2 (G) and 4 (H) days of DSS, normalized to the PV+ soma size (n = 107-175 cells from 4-6 mice/group and 63-93 cells from 3-4 mice/group for the 2- and 4-day timepoints, respectively). **(I)** Traces and mIPSC frequency from excitatory neurons in control and 4-day DSS (n = 12 cells; 3-4 mice/group). **(J)** Density of PV+ (Syt2+Gephyrin+) synapses on excitatory neurons (NeuN) at 4 days of DSS (n = 72-115 cells; 4-6 mice/group). Arrowheads: colocalization. **(K-L)** RAM labeling schematic (K) and density of GFP+ neurons in control and 4-day DSS mice (L); n = 5-7 mice/group. *p<0.05, **p<0.01, ***p<0.001, ****p<0.0001; unpaired t-test (B, C, I, L), one-way ANOVA with Tukey’s test (D), Kolmogorov-Smirnov test (G, H, J). Data: B, C, I, L, mean ± SEM; D, G, H, J, median and quartiles.

To identify which cellular processes drive the early PV+ response, we examined the identity of the DEGs from our snRNA-seq experiment. Upregulated genes encoded synaptic and cytoskeletal proteins, ion channels, and RNA-processing factors, along with Brevican (*17*), a key structural component of PNNs (Fig. 2E and table S2). Because PNNs constrain the capacity of PV+ interneurons for synaptic remodeling (*16*, *17*, *28–30*), we reasoned that they change alongside PV+ activity. Indeed, PNN volume around IC PV+ interneurons, labeled with Wisteria floribunda agglutinin (WFA), increased at day 2 of DSS, coinciding with peak PV+ activation (Fig. 2, F and G). By day 4, when PV+ activity had declined, PNNs decreased below control levels, without increased proteolytic Brevican cleavage (*31*) (Fig. 2, F and H, and fig. S5A), indicating a transition to a plasticity-permissive, low-PNN state.

We next asked whether the reduction in PV+ and SST+ activity at day 4 is accompanied by functional changes in synaptic inhibition of excitatory neurons. Whole-cell patch-clamp recordings from IC pyramidal neurons showed that the frequency of miniature inhibitory postsynaptic currents (mIPSCs) was reduced at 4 days of DSS (Fig. 2I), but unchanged at 2 days (fig. S5B). Spontaneous IPSCs followed the same time course (fig. S5C). Structural synapse analysis confirmed these functional observations: the number of PV+ presynaptic puncta (Syt2+) colocalizing with the postsynaptic protein Gephyrin and apposed to the soma of excitatory neurons was reduced at day 4 (Fig. 2J). Consistent with this disinhibition, activity-dependent labeling with the Robust Activity Marking (RAM) system (*32*) revealed an increased density of labeled neurons in the IC after 4 days of DSS compared to water-drinking controls (Fig. 2, K and L). RAM labeling was not significantly increased at a lower DSS dose (fig. S4J), matching the dose dependence of the PV+ response. Excitatory transmission onto pyramidal neurons was unchanged, as were the intrinsic electrical properties of both pyramidal neurons and PV+ interneurons (fig. S5, D to H). Together, these results establish that gut inflammation drives a severity-dependent, biphasic sequence of inhibitory circuit states in the IC: an initial high-PV, high-PNN configuration is followed by a plasticity-permissive low-inhibition state in which inhibitory output onto excitatory neurons is selectively reduced and IC neurons are recruited.

## VIP+ interneurons gate the transition of the IC to a disinhibited state

We next investigated what drives the IC from the initial high-PV+ state into the disinhibited configuration. Because microglia strip inhibitory synapses during peripheral inflammation (*33*, *34*), we first tested their contribution. After DSS, IC microglia showed no morphological signs of activation (Fig. 3A and fig. S6, A to C), did not upregulate disease-associated or inflammatory microglial genes (*35*) (fig. S6, D and E, and table S2), and did not engulf more PV+ presynaptic puncta (fig. S6F). We reasoned that if microglia contribute to shifting the network to a low PV+ state, their depletion should prevent the reduction in active PV+ cells at 4 days. We administered the CSF-1R inhibitor PLX5622 to deplete macrophages, including microglia, and then induced colitis by adding DSS. The reduction in the fraction of active PV+ cells after 4 days of DSS persisted in microglia-depleted mice (Fig. 3B). This result was not attributable to incomplete microglial depletion in the DSS context, nor to PLX5622 off-target effects on colitis severity (fig. S6, G to J). The reconfiguration therefore does not require microglia, pointing to a neuronal mechanism.

**Fig. 3.**
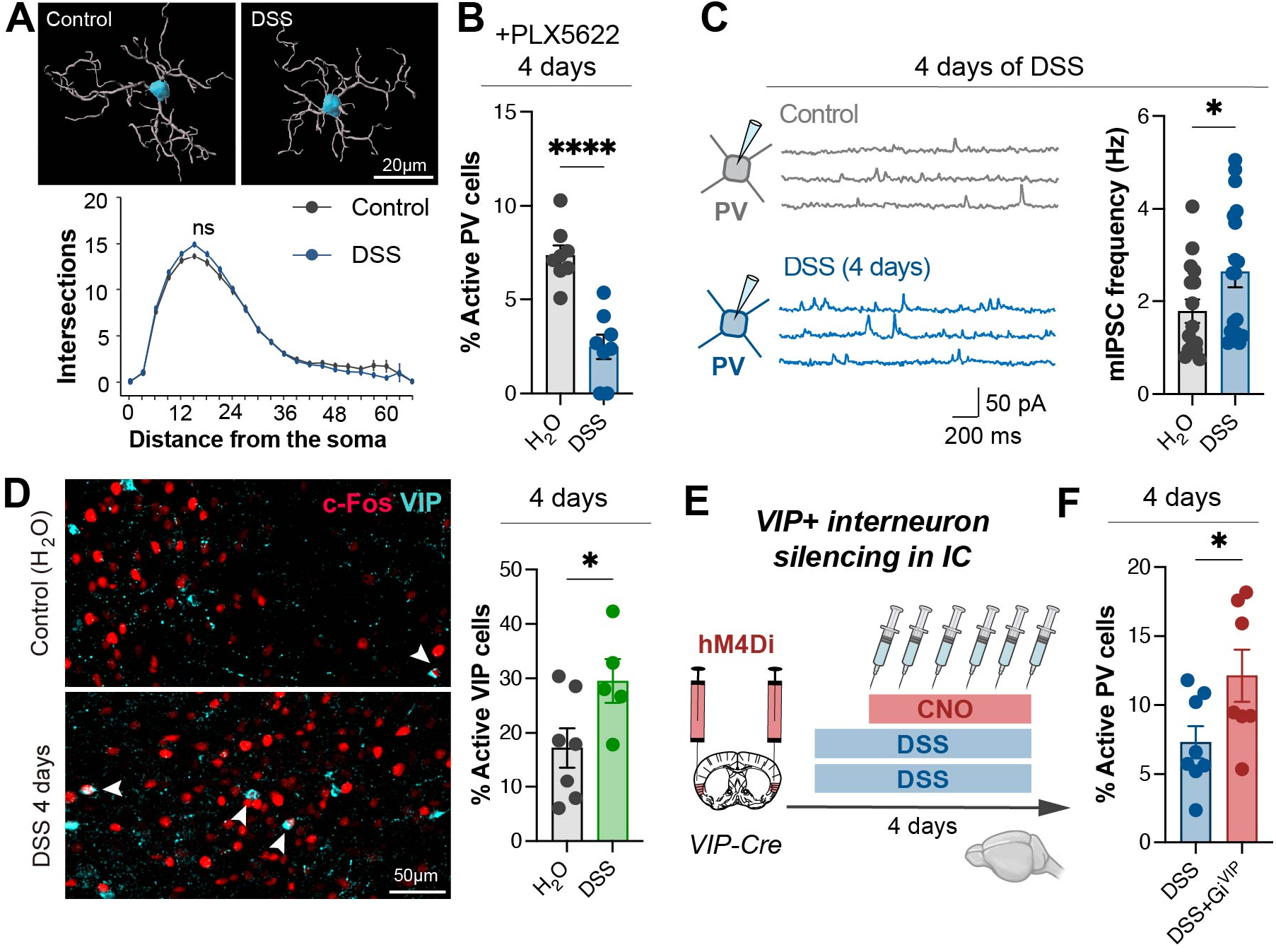
VIP+ interneurons, but not microglia, mediate the late suppression of IC inhibitory interneurons. **(A)** Representative skeletonized microglia and Sholl analysis in the IC at 4 days of DSS (n = 215 control cells and 221 DSS cells from 5 mice/group). **(B)** Fraction of c-Fos+ PV+ interneurons in the IC at 4 days of DSS in PLX5622-treated (PLX+) mice (n = 6-8 mice/group). **(C)** Recording schematic, representative traces and mIPSC frequency in S5E2-GFP+ PV+ interneurons at 4 days of DSS (controls: n = 16-20 cells from 3 mice; DSS: n = 18-19 cells from 5 mice). **(D)** Representative images and fraction of c-Fos+ VIP+ interneurons in the IC at 4 days of DSS (n = 5-7 mice/group). Arrowheads indicate c-Fos+ VIP+ interneurons. **(E-F)** Schematic of chemogenetic silencing of IC VIP+ interneurons during the day 2-4 window of DSS (E) and fraction of c-Fos+ PV+ interneurons in the IC at 4 days of DSS in control and VIP-Gi mice (F); n = 7-8 mice/group. *p<0.05, ****p<0.0001; linear mixed-effects model with random intercepts and slopes per cell (A), unpaired t-test (B, D, F), Mann-Whitney test (C). Data are mean ± SEM.

Disinhibitory circuits transiently suppress PV+ and SST+ output and open time-limited plasticity windows during learning (*16*, *18*, *19*, *28*). We therefore asked whether PV+ interneurons themselves receive more inhibition after DSS. Whole-cell recordings from S5E2-GFP-labeled PV+ interneurons in the IC (*36*) showed that the frequency of miniature inhibitory postsynaptic currents (mIPSCs) increased after 4 days of DSS compared to controls (Fig. 3C and fig. S7, A and B). PV+ interneurons receive inhibition from other PV+ cells, as well as SST+ and VIP+ interneurons (*37*, *38*). To identify the cellular source of the increased inhibition, we first labeled PV-PV synapses and found that they did not differ significantly from controls (fig. S7C). Because VIP+ interneurons gate the transition of PV+ cells to a high-plasticity state during learning (*16*, *28*), we next examined whether IC VIP+ interneurons are activated during immune encoding. The fraction of c-Fos+ VIP+ interneurons increased by day 4, with a non-significant trend already apparent at day 2 (Fig. 3D and fig. S7D). To test whether VIP+ activation is required for the low-inhibition transition induced by DSS, we silenced VIP+ interneurons using Cre-dependent inhibitory chemogenetics (hM4Di) in *VIP-Cre* mice starting at 24 hours of DSS (Fig. 3E). Consistent with the disinhibitory logic, silencing VIP+ interneurons prevented the day-4 fall in PV+ activity (Fig. 3F). VIP-mediated inhibition is therefore the neuronal source of the PV+ suppression that occurs in the IC days after gut inflammation.

## Trpv1+ sensory neurons drive IC inhibitory reconfiguration across immune challenges

Peripheral immune signals reach the brain through both humoral and neural routes (*39*). To determine whether IC PV+ interneurons are recruited through an ascending neural pathway, we first asked whether their activation involves increased synaptic input. At day 2 of DSS, the density of VGluT2+ excitatory synaptic puncta onto PV+ interneurons was increased (Fig. 4, A and B). To identify the afferents involved, we performed subdiaphragmatic vagotomy (SDV), which removes vagal innervation of the gut (Fig. 4C and fig. S8A). At 2 days of DSS, the increase in PV+ and VIP+ activation was absent in SDV mice (Fig. 4, D and E), although SDV alone raised the baseline activation of both populations. DSS-induced pathology was not altered by SDV (fig. S8B). Because SDV disrupts efferent as well as afferent fibers and can alter immune profiles through the cholinergic anti-inflammatory reflex (*40*, *41*), we employed a complementary genetic approach. We used *Trpv1-DTR* mice to selectively ablate Trpv1+ sensory neurons, a population that includes both spinal and vagal afferents innervating the gut (*42*, *43*). Daily diphtheria toxin (DT) injection for three weeks ablated Trpv1+ neurons, confirmed by the hot plate test and TRPV1 immunostaining (Fig. 4F and fig. S8, C and D). DT treatment alone did not alter baseline PV+ cell activation and, consistent with the vagotomy results, Trpv1-ablated mice showed no PV+ cell activation in response to DSS (Fig. 4G). Trpv1 ablation also reduced the VIP+ interneuron activation triggered by DSS (Fig. 4H), indicating that the same sensory population contributes to the recruitment of both interneuron types.

**Fig. 4.**
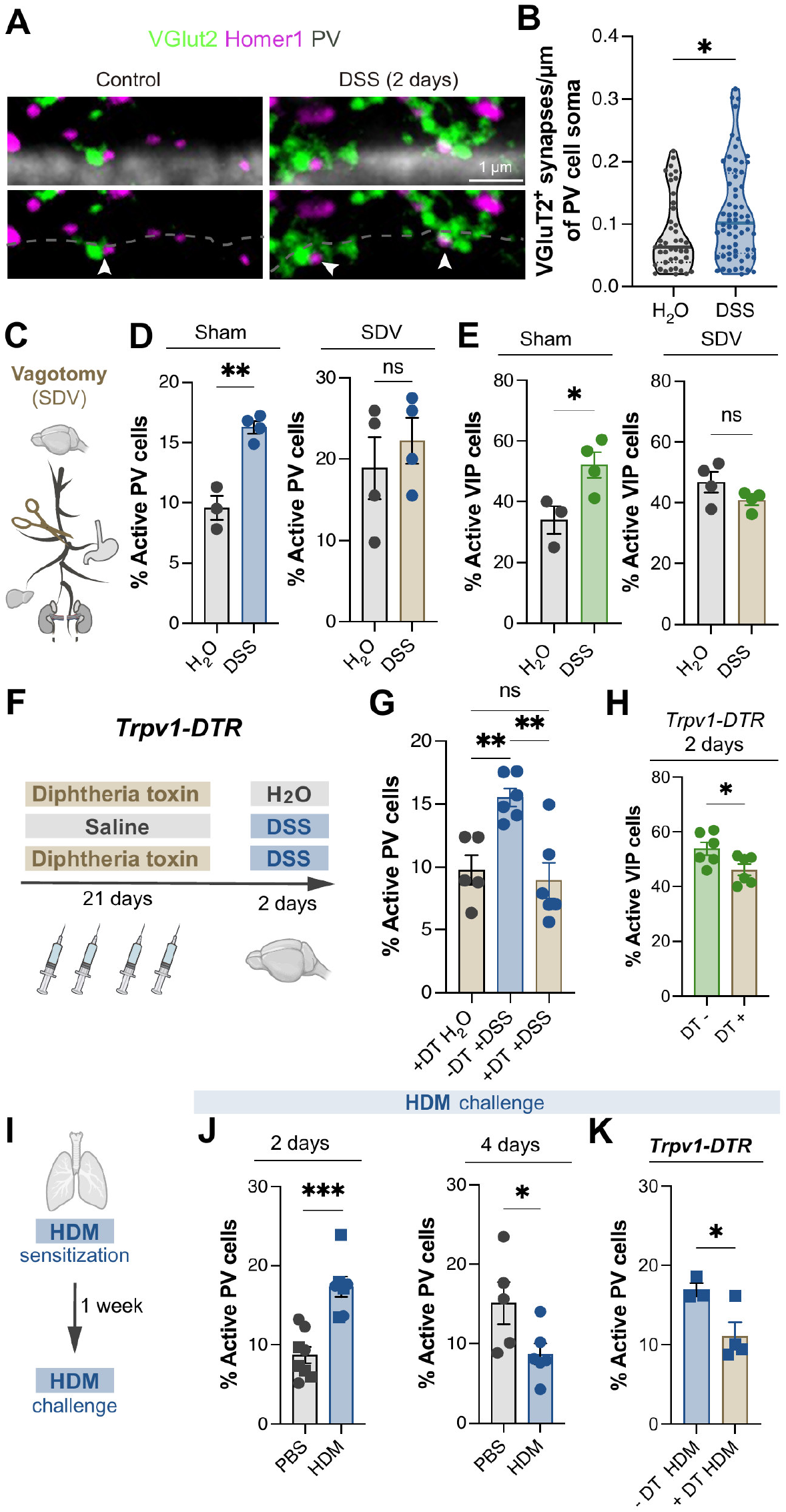
Vagotomy and Trpv1+ neuron ablation prevent IC PV+ and VIP+ interneuron activation after gut and airway inflammation. (**A-B**) Representative images (A) and density (B) of VGluT2+Homer1+ excitatory synapses on PV+ interneurons in controls and 2-day DSS (n = 43-64 cells; 5-6 mice/group). Arrowheads: synapses. **(C-D)** Subdiaphragmatic vagotomy (SDV) schematic (C) and fraction of c-Fos+PV+ interneurons in sham and SDV mice (D); n = 3-4 mice/group. **(E)** Fraction of c-Fos+VIP+ interneurons in sham and SDV mice; n = 3-4 mice/group. **(F-G)** Trpv1+ neuron ablation by diphtheria toxin (DT) in Trpv1-DTR mice (F) and fraction of c-Fos+PV+ interneurons in DT-treated controls, DSS without DT, and DSS+DT mice (G); n = 5-6 mice/group. **(H)** c-Fos+ VIP+ interneurons at 2 days of DSS in Trpv1-DTR mice ± DT; n = 6 mice/group. **(I-J)** HDM challenge schematic (I) and fraction of c-Fos+PV+ interneurons 2 and 4 days after HDM challenge (J); n = 7-8 (2 days) and 5-6 (4 days) mice/group. **(K)** Fraction of c-Fos+PV+ interneurons 2 days after HDM challenge in Trpv1-DTR mice ± DT; n = 3-4 mice/group. *p<0.05, **p<0.01, ***p<0.001; Kolmogorov-Smirnov test (B), unpaired t-test (D, E, H, J, K), one-way ANOVA with Tukey’s test (G). Data are mean ± SEM; in B, median and quartiles. Circles and squares indicate male and female mice, respectively.

To determine whether this pathway operates beyond gut-derived signals, we used house dust mite (HDM)-induced airway allergic inflammation (*44*), which targets the airways and engages distinct inflammatory mediators and effector immune cells (Fig. 4I and fig. S9). PV+ interneurons were activated 2 days after HDM challenge in both males and females (Fig. 4J and fig. S10, A and B). At 4 days after challenge, PV+ activity was suppressed in males but not in females (Fig. 4J and fig. S10C). HDM sensitization, which elicits a weaker peripheral immune response than challenge (fig. S9, B and C), triggered the first phase of PV+ response but not the second (fig. S10, D and E). As after DSS, Trpv1+ neuron ablation abolished the PV+ activation induced by HDM (Fig. 4K and fig. S10F). Together, these results demonstrate that Trpv1+ sensory neurons are required for the recruitment of IC inhibitory interneurons across organs and types of immune response. An independent study reached the same conclusion, showing that Trpv1+ neuron ablation or systemic analgesia during peritoneal inflammation prevents the IC from forming a retrievable immune representation (Megan Sammons & Asya Rolls, personal communication).

## Disrupting inhibitory circuit reconfiguration impairs immune encoding in the IC

Having established how peripheral immune challenges reconfigure IC inhibitory circuits, we asked whether this reconfiguration is causally required for immune engram formation. To test this, we first established the immune engram paradigm (*4*) in our experimental conditions. We injected the mid-posterior IC of *FosTRAP2* mice (*45*, *46*) with AAVs encoding a Cre-dependent excitatory chemogenetic actuator (hM3Dq) together with a fluorescent reporter, and administered tamoxifen 48 hours after DSS onset to permanently label activated ensembles (Fig. 5A). Because tamoxifen co-administration worsened colitis, we used 3.5% DSS in these experiments to achieve pathology equivalent to that of 5% DSS (fig. S11A). One month later, chemogenetic reactivation of these ensembles with clozapine-N-oxide (CNO) reinstated immune cell infiltration in the colon mucosa (Fig. 5, B and C).

**Fig. 5.**
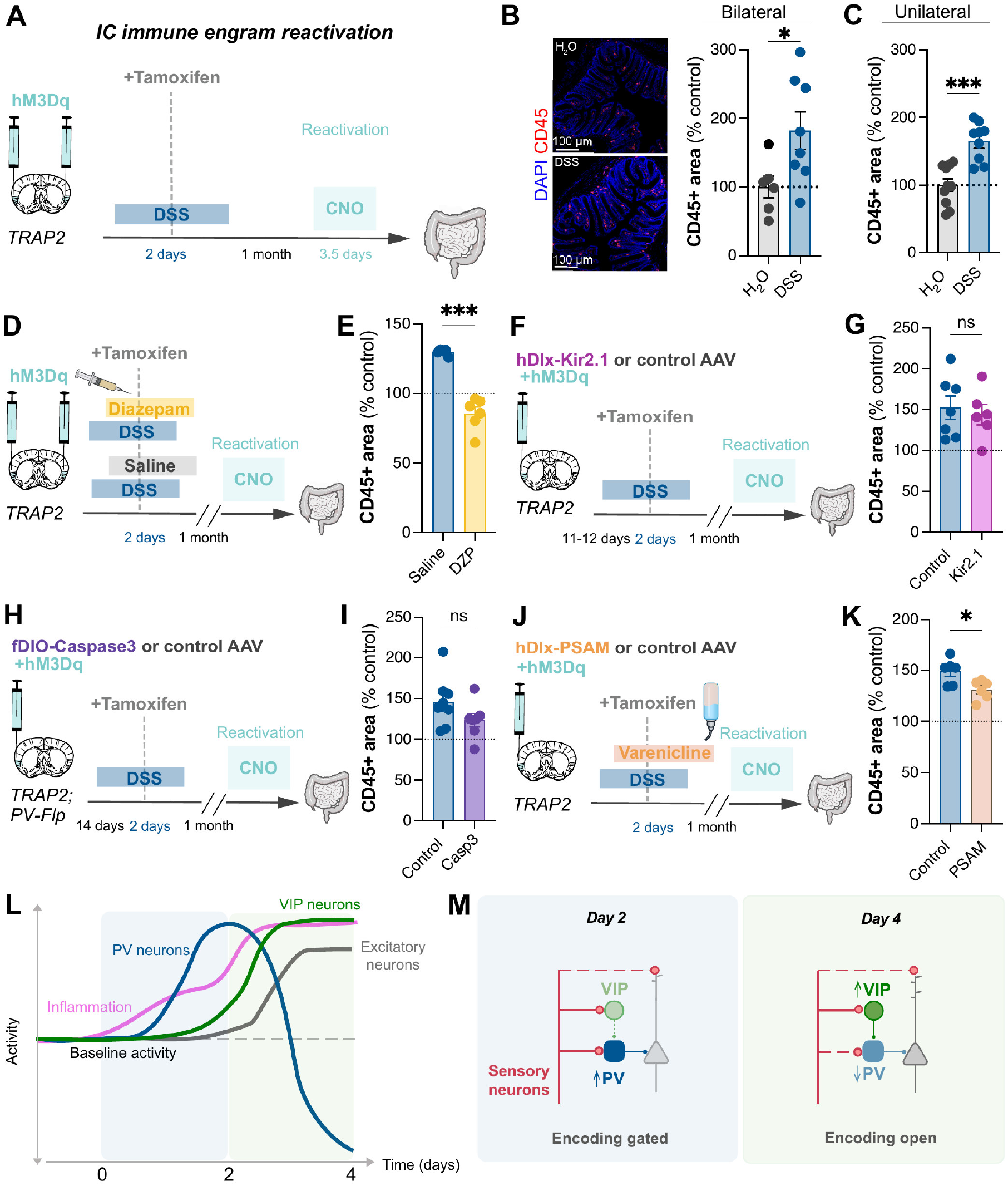
Inhibitory circuit reconfiguration during the encoding window is required for immune engram formation. (**A-C**) Schematic of the immune engram reactivation experiment (A) and representative images of CD45+ cell infiltration in the colon mucosa and quantification after bilateral (B) or unilateral (C) reactivation of IC TRAPed neurons; n = 6-10 mice/group. **(D-E)** Schematic (D) and CD45+ area after reactivation in saline- and diazepam (DZP)-treated mice (E); n = 5-6 mice/group. **(F-G)** Chronic interneuron silencing with hDlx-Kir2.1 (F) and CD45+ area after reactivation in control and hDlx-Kir2.1 mice (G); n = 6-7 mice/group. **(H-I)** PV+ interneuron ablation in *TRAP2;PV-FlpO* mice (H) and CD45+ area after reactivation in control and fDIO-Caspase3 mice (I); n = 7-8 mice/group. **(J-K)** Dual chemogenetic experiment with hDlx-PSAM (J) and CD45+ area after reactivation in control and hDlx-PSAM mice (K); n = 6 mice/group. **(L)** Time course of peripheral inflammation and IC cell-type activity. **(M)** Working model of the IC inhibitory circuit and ascending input. On day 2, sensory drive engages PV+ interneurons and gates encoding. By day 4, the same afferents recruit VIP+ interneurons, which suppress PV+ interneurons and open the plasticity window. Sensory input is drawn as direct but is likely polysynaptic. *p<0.05, ***p<0.001; unpaired t-test (B, C, G, I, K), Welch’s t-test (E). Data are mean ± SEM.

If the transition to a disinhibited state is required for encoding, maintaining elevated inhibitory tone through the window should prevent immune engram formation. We tested this by administering diazepam, a systemic potentiator of GABA_A_ receptors, from day 2 of DSS, together with tamoxifen to label immune-activated neurons (Fig. 5D). Reactivation of DSS-TRAPed neurons one month later failed to reinduce colon inflammation in diazepam-treated mice (Fig. 5E and fig. S11B). We then asked whether this reflects a general sensitivity to perturbation of IC inhibition. Chronic suppression of inhibitory interneurons with Kir2.1 driven by the interneuron-selective hDlx enhancer (*47*, *48*), or ablation of PV+ interneurons with Caspase3 before DSS, did not impair immune engram reactivation (Fig. 5, F to I; fig. S11, C to E). Mice consumed comparable volumes of drinking solution across all manipulations (fig. S11D).

Because diazepam acts on all GABA_A_ receptors, including those on immune cells (*49*, *50*), we next tested whether activating IC interneurons alone is sufficient to block encoding. This required manipulating two neuronal populations independently within the same circuit, which we achieved using orthogonal chemogenetic actuators driven by different ligands at different times. Because unilateral reactivation was as effective as bilateral reactivation (Fig. 5C), we unilaterally co-injected TRAP2 mice with a Cre-dependent hM3Dq for later reactivation and, in one group, an AAV in which the hDlx enhancer (*47*, *48*) drives expression of the chemogenetic actuator PSAM4-5HT3 (Fig. 5J and fig. S11F). Mice received tamoxifen during DSS to label immune-activated neurons, together with varenicline in the drinking water to activate PSAM4-expressing interneurons across days 1-4. Varenicline did not alter colitis severity, and mice consumed comparable volumes of drinking solution (fig. S11, G to I). One month later, CNO-triggered reactivation failed to reinduce colon inflammation in mice whose IC interneurons had been held active during the encoding window (Fig. 5K), reproducing the diazepam result with regional and cell-type specificity. Together, these results establish that inhibitory circuit reconfiguration during the encoding window is required for immune engram formation in the IC (Fig. 5, L and M).

## Discussion

A challenge the brain faces when encoding immune experiences is that they develop gradually over several days. Here, we find that Trpv1+ sensory neurons drive a coordinated, multi-day reconfiguration of the local inhibitory network in the IC, which is required for immune engram formation. These results establish a body-to-brain pathway that converts immune responses into cortical engrams.

Disinhibitory circuit motifs are a recurring solution the cortex uses to encode experience (*23*, *51*). However, the stimuli that engage these motifs are typically brief, whereas peripheral immune challenges persist for days. The ordered reconfiguration we observe provides a mechanism by which the cortex adapts the timing of plasticity to the slow timescale of a peripheral physiological signal. The early high-PV phase may act as a filter that raises the threshold for recruitment of excitatory neurons, restricting engram allocation to sustained or suprathreshold inputs, a form of gain control (*52*, *53*). Consistent with this, weaker challenges such as low-dose DSS or HDM sensitization elicit the early PV+ response but not the subsequent transition to disinhibition, and immune engram formation is insensitive to lowering the basal state of the inhibitory network by overexpressing Kir2.1 in inhibitory neurons. Consistent with the severity dependence of the transition we describe, nociceptive drive scales the strength and composition of immune recall (Megan Sammons & Asya Rolls, personal communication). Alternatively, the initial PV+ activation may be important to establish engram sparsity (*12–14*). Whichever function the early phase serves, the VIP-gated transition to a disinhibited state opens the encoding window, paralleling the classical framework of engram formation (*12–14*, *16*).

The delay in this transition could reflect weaker or more indirect afferent recruitment of VIP+ than PV+ neurons (*54*, *55*). Because VIP+ activation also requires Trpv1+ sensory neurons, both phases originate from the same peripheral afferents, and the delay therefore arises downstream of a common ascending signal. Consistent with this, PV+ interneurons are enriched for AMPA receptor subunits that support rapid, high-efficacy transmission, whereas the class containing VIP+ cells (Adarb2+) relies more on NMDA and metabotropic signaling, which may require stronger or more sustained input (fig. S7E). These dynamics determine when excitatory neurons can be recruited, whereas which neurons are allocated to the engram is likely set by the coincidence of immune-driven excitatory drive with the disinhibited window.

Dynamic PNN remodeling accompanies the biphasic circuit response and may provide a structural substrate for its temporal gating. PNN accumulation during the high-PV phase would constrain synaptic plasticity and protect the circuit from premature disinhibition (*56*, *57*), while PNN loss at day 4 would remove this constraint and permit the structural remodeling required for engram recruitment. A key question is what triggers PNN changes. PV+ interneurons can cell-autonomously reduce PNNs, although the relationship is bidirectional (*16*, *17*, *58*). Because extracellular matrix remodeling is a hallmark of inflammation (*59*, *60*), an intriguing possibility is that peripheral immune signals also regulate the expression of PNN components directly. Consistent with this, we found that Brevican and Aggrecan promoters contain predicted JAK-STAT binding sites, and astrocytes, the principal source of PNN components, are the most transcriptionally responsive non-neuronal cell type in our snRNA-seq dataset (fig. S6D).

Gain control may also operate upstream of the cortex. Trpv1+ neurons convey noxious and irritant stimuli, and they may act as salience signals for immune encoding. Consistent with a salience function, augmenting nociceptive drive during inflammation strengthens subsequent immune recall and shifts its cellular composition, whereas analgesia abolishes it (Megan Sammons & Asya Rolls, personal communication). That the requirement for Trpv1+ sensory neurons holds across gut and airway challenges, which engage distinct afferent populations, suggests that these afferents convey both organ-specific information and a shared salience signal. The organ-specific component would determine what is encoded, while the shared salience component would determine whether encoding occurs at all. Spinal sensory neurons terminate in the dorsal horn and are likely to relay immune information to the brain through spinothalamic pathways (*61*). Vagal sensory neurons terminate in the nucleus of the solitary tract, and immune information likely reaches the IC through downstream thalamic and parabrachial nodes (*62*, *63*). Neurons in the thalamus and parabrachial nucleus have VGluT2+ terminals, consistent with the increased VGluT2+ input onto IC PV+ interneurons during DSS.

The circuit we describe is engaged each time the body mounts an immune response. Inflammatory bowel disease, asthma, and other relapsing inflammatory conditions are characterized by symptoms that recur after resolution of the inciting event (*64*). We propose that repeated immune challenges may progressively stabilize a disinhibited state in the IC, lowering the threshold for immune engram formation and retrieval, such that the cortex itself amplifies and sustains peripheral inflammation. Because encoding depends on a time-limited disinhibitory window, that window is in principle a therapeutic target, and benzodiazepines blocked immune engram formation in our experiments. Instead of broadly suppressing peripheral immunity, stabilizing IC inhibitory tone during immune responses may represent a strategy to modulate immune engram formation before it drives disease chronicity.

## Acknowledgments

The authors thank Drs. S. Tomita and M. Higley for critical reading of the manuscript, Drs. Rui Chang and Quancheng Zhao for vagotomy assistance, the Gracheva lab for fUSI equipment, Drs. Isaac Chiu and Mark Hoon for *TRPV1-DTR* mice, the Sestan lab for assistance with snRNA-seq preparation, Dr. Renato Frischknecht for Brevican antibodies, and Dr. D. Camacho for initial assistance with the HDM model; I. Sun, A. Singh, and K. Carrier for contributing to the quantification of figs. S7D, S4D, and S10A, respectively. Illustrations in Figs. 2K, 4I, and 5A are from NIAID NIH BioArt Source.

## Funding

Charles H. Hood Foundation Child Health Award (EF)

Brain and Behavior Research Foundation Young Investigator Grant 31507 (EF) Esther A. and Joseph Klingenstein Fund (EF)

Ludwig Pilot Grant Program Award, Carol and Gene Ludwig Program for the Study of Neuroimmune Interactions in Dementia at Yale University (EF)

National Institutes of Health grant R01AG098889 (EF) National Institutes of Health grant R01MH142612 (EF)

Thomas Kingsley Lawrence ’19 Program in Brain Research (EF) Kenneth Rainin Foundation Innovator Award (EF)

Arnold and Mabel Beckman Foundation Beckman Young Investigator Award (EF)

Alzheimer’s Disease Research Center Scholar Award, National Institutes of Health grant P30AG066508 (EF)

Wu Tsai Institute startup funds (EF) Rudolf J. Anderson Fellowship (EF, ZHG)

National Institutes of Health grant P20GM148302, CNDD Genomics and Bioinformatics Core at the Medical University of South Carolina (SS, SB)

## Author contributions

Conceptualization: EF

Methodology: ZHG, ST, EF

Software: ZHG, SS, SB, EF

Validation: ZHG, EF

Formal analysis: ZHG, ST, MW, CL, SYF, PCD, EKF, SS, AI, SB, AC, EF

Investigation: ZHG, ST, MW, CL, AW, SYF, PCD, EKF

Resources: SB, AC, EF

Data curation: SS

Writing – original draft: EF

Writing – review & editing: ZHG, ST, MW, CL, SYF, EKF, SS, AW, PCD, AI, SB, AC, EF

Visualization: ZHG, EF

Supervision: EF

Project administration: EF Funding acquisition: EF

## Competing interests

Authors declare that they have no competing interests.

## Data, code, and materials availability

Single-nucleus RNA sequencing data have been deposited in NCBI GEO under accession number GSE329920. Analysis code is available at https://github.com/BioinformaticsMUSC/FavuzziEtAl_snRNA_mInsularCortex, and an interactive browser is available at http://bioinformatics-musc.shinyapps.io/Favuzzi_snRNA_mm_InsCortex_Preliminary. All plasmids generated in this study are available from the corresponding author on request. All other data are available in the main text or the supplementary materials.

Supplementary Materials

Materials and Methods

Figs. S1 to S11

Tables S1 and S2

References (65–82)

Data S1

## Materials and Methods

### Mice

The following mouse lines were obtained from the Jackson Laboratory: C57BL/6J (“WT”, strain # 000664), Fostm2.1*^(icre/ERT2)Luo^*/J (“*TRAP2*”, strain # 030323), B6.Cg-*Pvalb^tm4.1(flpo)Hze^*/J (“*PV-FlpO*”, strain # 022730), *Vip^tm1(cre)Zjh^*/J (“*Vip-Cre*”, strain # 010908), B6.Cg-*Gt(ROSA)26Sor^tm75.1(CAG-tdTomato*)Hze^*/J (“*Ai75D*”, strain # 025106). *Trpv1-DTR* (*42*) mice were generously provided by Drs. Mark Hoon and Isaac Chiu. All lines were maintained on a C57BL/6J background and genotyped by PCR from ear-biopsy genomic DNA. Experimental animals were heterozygous for all alleles above.

All animal husbandry and experimental procedures were conducted in accordance with protocols approved by the Yale University Institutional Animal Care and Use Committee (IACUC; protocol 2025-20465) and with National Institutes of Health (NIH) guidelines. All efforts were made to minimize animal pain and reduce the number of animals used. Mice were group-housed under a standard 12:12-hour light-dark cycle with ad libitum access to food and water, except where a specialized diet is specified below. Adult mice 8-12 weeks of age at the start of the experiment were used. Male mice were used for DSS experiments (except in fig. S4 and in the sn-RNAseq experiment, where both sexes were used). Both males and females were used for the HDM experiments, with the sex of the animals indicated in each figure legend. Some tissue was used for more than one analysis; each such use is indicated in the relevant figure legend, which also references every earlier panel that drew on tissue from overlapping animals. Whenever possible, assignment of mice to experimental groups was randomized (e.g., littermates assigned to PBS or HDM groups). To minimize variability, littermates were used in all experiments involving a comparison between conditions and often included the use of multiple litters. Brains from the two conditions were always processed, imaged and analyzed together. When mice from multiple litters were pooled, they were collected simultaneously or within a short period of time (less than a month) and each collection always included both experimental and control groups. All main phenotypes were confirmed in independent cohorts, and analyses were performed blind to treatment group.

### DSS model of acute colitis

We administered dextran sulfate sodium (DSS) to model acute colonic inflammation with features of ulcerative colitis in mice (*65*) and examine the effects of colon inflammation on the brain. Across experiments, fresh feces were inspected daily and assigned an ordinal “feces pathology score” on a five-point severity scale (0, normal; 1, change in color; 2, soft feces; 3, trace blood; 4, frank blood), so that higher scores denote more severe disease. In experiments using WT mice, access to drinking water was replaced with 5% (w/v) DSS (MP Biomedicals, catalog # 160110, Lot #s U1122304862-1, U1122304862-4, and G1124001641A) in sterile drinking water. Water bottles were weighed to confirm comparable intake between groups, and intake did not differ between control and DSS mice. The DSS solution was replaced every 3 days. The experiments in figs. S1C and S4, E and F, used 3% DSS to model milder colon inflammation. In experiments involving tamoxifen administration, 3.5% DSS was used because 5% DSS caused mortality within a week of tamoxifen treatment; 3.5% DSS produced equivalent pathology (fig. S11A). Colitis pathology and brain responses were comparable across DSS lots. Across experiments, DSS administration began at approximately 19:00, at the onset of the dark phase, so that day 1 of the model corresponds to the first full dark period of exposure. Control animals received identical sterile water without DSS added. At perfusion fixation and tissue extraction (described below), colon length was measured from the cecum to the anterior pubic symphysis, with care not to stretch the tissue, as an additional index of colitis severity. Spleen length was measured in the same way in the microglia-depletion experiment.

### Functional ultrasound imaging (fUSI)

#### Animal preparation

Resting-state fUSI data were acquired transcranially in lightly sedated mice. The scalp was shaved with an electric trimmer, mice were induced with 4% isoflurane and placed in a stereotaxic frame, and anesthesia was then maintained with a combined ketamine/xylazine-isoflurane regimen: ketamine (100 mg/kg) and xylazine (10 mg/kg body weight) were given intraperitoneally (i.p.), and isoflurane (0.2-0.5%) was delivered by inhalation for the remainder of the session. Ophthalmic gel was applied to prevent corneal drying. Body temperature was maintained at 37 ± 0.5°C using a temperature controller (Physitemp TCAT-2). Respiratory rate (130-150 breaths/min) was manually monitored. Ultrasound gel (Aquasonic, Parker) was liberally applied in a ∼3 mm layer to the scalp with a 10 mL syringe, completely covering the imaging window. The ultrasonic probe was positioned approximately 1 mm above the skull using an XYZ motorized controller (Iconeus, Paris, France). The same mice were imaged at all three time points, and sessions were separated by at least 48 h.

#### fUSI acquisition

Functional ultrasound imaging was performed using an Iconeus One scanner (Iconeus, Paris, France) equipped with a 15-MHz multiarray probe (256 channels) mounted on a four-axis motorized stage. Multi-slice imaging was achieved by rapid motorized movement of the probe between slices. To ensure sufficient temporal resolution for resting-state analyses, data were acquired at a sampling frequency of 0.4 Hz (1 data point every 2.5 seconds) in two contiguous coronal imaging stacks (an anterior and a posterior stack). The posterior stack comprised three coronal planes spaced 0.60 mm apart, centered at AP −2.06, −1.46 and −0.94 mm relative to Bregma. The anterior stack comprised four coronal planes spaced 0.72 mm apart, centered at AP −0.94, −0.22, +0.50 and +1.18 mm relative to Bregma. Imaging was performed with the IcoScan software and each imaging session began with an angiographic scan covering the brain (Angio 3D option). The coordinates were then entered in IcoScan, corresponding to the probe position, allowing the targeted plane to be imaged. Probe positioning was verified using a vascular landmark defined by the bifurcation of the internal carotid artery into the anterior choroidal artery. Functional connectivity was measured in the same mice at three time points: baseline, Day 2 and Day 4 of DSS (or water in controls). Each resting-state acquisition lasted 5 min (5 for the posterior recording and another 5 for the anterior recording). The probe provided a field of view 14.1 mm wide and approximately 1.5 cm deep. Power Doppler images, which report relative cerebral blood volume, were reconstructed from 11 tilted plane-wave compoundings per frame at a pulse repetition frequency of 5.5 kHz, giving an in-plane resolution of 100 × 100 μm. Compounded frames were acquired at 0.2s intervals, and each plane was sampled every 2.5s as the probe cycled through the stack.

#### fUSI data analysis

Acquisitions were loaded into IcoStudio and regions of interest were selected from the “functional connectivity” menu. Frames displaying strong motion artifacts were rejected using the built-in motion-detection tool in IcoStudio. Matrices of the mean Pearson’s correlation coefficient between pairs of brain regions were exported for each animal and in each imaging stack using the IcoStudio software (table S1). Anatomically defined regions of interest (ROIs) were taken from the Allen Mouse Brain Atlas ontology and were selected to give broad anatomical and imaging coverage, guided by previous human and rodent studies (*4*, *66*). Functional connectivity (FC) between each pair of ROIs was computed as the Pearson correlation between their mean regional power-Doppler time courses, yielding one region-by-region FC matrix per mouse, stack and time point; these matrices were the input to all subsequent group and time comparisons. Analyses were restricted to the a priori ROI selection for each volume (44 ROIs in the anterior stack and 54 in the posterior stack), with the left and right hemispheres kept as separate ROIs. Gustatory areas were excluded from the posterior selection because they were not reliably captured in that stack. The insular cortex (IC) was defined as the agranular insular area (left and right); a broader definition additionally including the adjacent visceral and gustatory areas gave equivalent results (data not shown). Correlation coefficients were Fisher z-transformed (arctanh) to stabilize their variance and better approximate normality before statistical testing.

#### fUSI statistical testing

Given the exploratory nature of this dataset and the small sample size, edge-level and region-level screens are reported with uncorrected p-values and are interpreted descriptively. The primary insular claims rest on single, pre-specified summary indices (insular nodal strength, its rate of change, and its increase relative to the global network change), which do not incur a multiple-comparison penalty. We used Claude Fable 5 (Anthropic) to write the code for the functional connectivity analyses and to generate the fUSI visualizations, based on analysis specifications and figure requirements provided by the authors. We checked all code and verified its outputs before use. fUSI analyses were performed in Python 3.10.12 using NumPy 2.2.6, pandas 2.3.3, SciPy 1.15.3, statsmodels 0.14.6 (linear mixed-effects models) and pingouin 0.6.1 (mixed-design and repeated-measures ANOVAs). Data were read and written with openpyxl 3.1.5, and figures generated with Matplotlib 3.10.9 and seaborn 0.13.2. To generate the group-averaged connectivity matrices for each group, volume, and timepoint, a group-average FC matrix was obtained by averaging the per-mouse Fisher z values across mice and converting back to the correlation scale (r) for display. The change in connectivity from baseline to Day 4 (Δr) was computed per group as the difference of the group-average correlation matrices at the two time points. To detect pairs that changed significantly from baseline within the DSS group, a paired t-test was run on the Fisher z change. Insula-to-network connectivity was defined, for each mouse and timepoint, as the mean Fisher z-transformed correlation of the left and right agranular IC seeds with all other imaged regions (the insular nodal strength). Recruitment was tested within the DSS group as the change from baseline to Day 4 with a two-tailed paired t-test. Stability in controls was assessed with a within-H_2_O repeated-measures ANOVA across days, and baseline equivalence with an independent-samples t-test (H_2_O vs DSS at baseline). Specificity to DSS was tested in two complementary, baseline-invariant ways, which yielded similar results: (i) a permutation test on the per-mouse baseline-corrected change scores (Day 4 minus baseline), comparing DSS with H_2_O (20,000 label shuffles, not shown); and (ii) a linear mixed-effects model of the insular index with a group-by-time interaction (time as a continuous variable, random intercept per mouse), where the interaction estimates the difference in the rate of change (slope) between groups. Per-mouse slopes were calculated as the change in Fisher z per day, from an ordinary least-squares fit across baseline, Day 2 and Day 4. To identify the most recruited regions without pre-selecting the insula ranking, every ROI was ranked by the change in its nodal strength (mean Fisher z-transformed correlation of that region with all other regions) from baseline to Day 4 in DSS mice. The whole-network average change was computed across all ROIs, and each region’s change was expressed as a fold of that average. For the edge-level analyses, for every ROI pair, the within-DSS change from baseline to Day 4 was tested with a paired t-test on the Fisher z change across DSS mice. To test whether the insular increase exceeded a network-wide change, we computed for each mouse, timepoint and volume a global connectivity index, defined as the mean Fisher z-transformed correlation across all region pairs not involving either agranular insular seed. The change from baseline to Day 4 was taken for the insular and the global index, and their per-mouse difference (Δinsula - Δglobal) was tested against zero with a two-tailed one-sample t-test, separately for each group and volume.

### Single-nucleus RNA sequencing

Mice were exposed to 5% DSS for 48 h prior to tissue extraction. We used 4 control mice (H_2_O, 2 females and 2 males) and 4 DSS mice (2 males and 2 females). Tissue from 4 mice was pooled per library, and 1 library was prepared per condition. For tissue extraction, mice were briefly anesthetized with isoflurane and acutely injected i.p. with Euthasol Solution. Animals were then rapidly exsanguinated through intracardiac perfusion with 25mL ice-cold (∼4°C) Hanks’ Balanced Salt Solution (HBSS; Corning, catalog # 21-022-CV). Following exsanguination, brains were rapidly removed and the right IC was dissected on ice while remaining submerged in ice-cold HBSS and with the assistance of a dissecting microscope. Prior to the first extraction and between animals, all tools and surfaces were decontaminated with “RNaseZap” RNase Decontamination Solution (Invitrogen, catalog # AM9780). Extracted insular cortex tissue was then rapidly frozen in a pre-chilled tube on dry ice. Tissue was then stored at -80°C until nuclei isolation. Nuclei isolation was performed according to a previously published protocol (*67*) with minor modifications. Specifically, we excluded the initial liquid nitrogen step and subsequently submitted samples to the Yale Center for Genome Analysis (YCGA) for library preparation and sequencing.

#### Construction and sequencing of 10x Genomics GEM-X Single Cell 3′ libraries

Library preparation and sequencing were performed by the Yale Center for Genome Analysis (YCGA) according to the manufacturer’s protocol (10x Genomics Chromium GEM-X Single Cell 3′ Reagent Kits v4). Briefly, the 2 samples were provided at 1280 nuclei per uL (H_2_O) and 1227 nuclei per uL (DSS). 13.8 µL of nuclei suspension with 29.5 µL of water were run to target capture of 10k nuclei per library. This resulted in approximately 17664 and 16933 nuclei being loaded for H_2_O and DSS libraries, respectively. Loaded nuclei were partitioned into nanoliter-scale gel bead-in-emulsions in which each barcoded gel bead carries an Illumina R1 sequence, a 16 nt 10x barcode, a 12 nt unique molecular identifier, and a 30 nt poly(dT) primer, so that reverse transcription within the emulsion yields barcoded, full-length cDNA from polyadenylated transcripts. cDNA was purified on silane magnetic beads, amplified by PCR, enzymatically fragmented and size-selected, and converted into Illumina paired-end libraries by end repair, A-tailing, adaptor ligation, and indexing PCR. Libraries were normalized to 2.0 nM, spiked with 0.3% bacteriophage PhiX control library to monitor run quality, and sequenced on an Illumina NovaSeq to a target of 200 million passing-filter clusters per sample and a minimum of 20,000 read pairs per nucleus, using 10 bp dual indices with 28 bp for Read 1 (16 bp barcode plus 12 bp unique molecular identifier) and 91 bp for Read 2 (cDNA insert). Base calls were transferred to the YCGA high-performance computing cluster and processed as described below.

#### Data processing and analysis

Raw base call (BCL) files were analyzed using CellRanger (v7.0.0) (*68*). The “mkfastq” command was used to generate FASTQ files and the “count” command was used to generate raw gene – cell expression matrices. Ambient RNA contamination was inferred and removed using CellBender (v0.2.0) with standard parameters. Mouse genome mm10 was used for the alignment, and GENCODE vM25 was used for gene annotation and coordinates (*69*).

Downstream analysis was performed in R with Seurat (v4.3.0) (*70*) and customized R scripts. Data from different samples (DSS and Ctrl) were merged into a unique single-cell object. Filtering was conducted by retaining nuclei that had unique molecular identifiers (UMIs) less than 25000, mitochondrial content less than 5 percent, number of Genes(nGene) greater than 250 and nuclear fraction(nf) greater than 0.5. Doublets were removed using scDblFinder (v1.12.0) (*71*) for a resultant expression matrix with 21,849 genes and 15644 cells. The SCTransform workflow was used for count normalization, initial integration, and to identify highly variable genes (*72*) using 30 principal components and a resolution of 0.5 for Louvain clustering and UMAP. Cluster marker genes were identified using FindAllMarkers using Wilcoxon Rank Sum test with the standard parameters. We identified 23 final distinct clusters, of which D1/D2 STR showing high expression in the striatum were removed. Cell annotation was done by Fisher enrichment analysis with Allen PFC brain data, reference mapping with Azimuth mouse motor cortex, and manual curation of known canonical markers. Genes differentially expressed were calculated between DSS and Ctrl. The R package LIBRA (v1.0.0) was used to perform zero-inflated regression analysis (*73*). Genes were defined as significantly differentially expressed at Benjamini-Hochberg correction FDR < 0.05 and abs (log2(Fold Change)) > 0.3. Intercellular communication network analysis was performed using the standard workflow of the R package ‘CellChat’ (v2.1.2) (*74*). Ligand-receptor analysis framework. LIANA (v0.1.12), based on the consensus of several interaction analysis methods and resources were used to detect interactions between cells. MultiNicheNet (https://github.com/saeyslab/multinichenetr) package was used to find the differences in communication between the samples. For Figure S3E and S3F, we used Claude Opus 5 (Anthropic) to overlay two separate graphs into a single plot. We confirmed that the combined plot matched the source graphs. Data are deposited in the Gene Expression Omnibus under accession GSE329920, and analysis code can be found at https://github.com/BioinformaticsMUSC/FavuzziEtAl_snRNA_mInsularCortex.git. Interactive sequencing data are available through ShinyCellPlus http://bioinformatics-musc.shinyapps.io/Favuzzi_snRNA_mm_InsCortex_Preliminary.

### Brain and colon tissue preparation for immunohistochemistry

Before perfusion fixation of tissues for postmortem immunohistochemistry, animals were deeply anesthetized with 0.03 mL of Euthasol (Virbac, VINV-CIII-0001) by i.p. injection. Animals were then transcardially perfused with 25 mL of PBS followed by 25 mL of 4% paraformaldehyde (PFA) in PBS. Brains were extracted intact from the skull and post-fixed in 4% PFA for 3 h at 4 °C, then cryoprotected in 30% sucrose in PBS at 4 °C for approximately 2-5 days. Colons were dissected between the cecum and the anterior pubic symphysis, standardizing the anatomical landmarks used for colon length measurement and for histological analysis of the distal colon, with care not to stretch the tissue. Colons were then post-fixed in 4% PFA for 48 h at 4 °C, followed by cryoprotection in 30% sucrose in PBS for approximately 24 h at 4 °C. A subset of colon tissue was transferred to 70% ethanol instead of 30% sucrose after postfixation and sent to Histowiz for “Wet-to-Digital H&E” service. Both brains and colons were sectioned on a sliding microtome (Leica, SM2010 R). Tissue was frozen and kept frozen on dry ice, thinly encased in 30% sucrose in PBS to maintain rigidity during sectioning. Brain tissue was sectioned coronally at 40 µm (collected in 5 series) and distal colon tissue was sectioned transversely at 80 µm along the proximo-distal axis. Tissue was stored in a cryoprotective solution (4 parts 1X PBS, 3 parts glycerol, and 3 parts ethylene glycol) at -20°C until immunohistochemistry was performed.

### Histowiz H&E and histopathology scoring

Colon tissue samples from control and DSS mice were placed in 70% ethanol and subsequently processed, paraffin embedded, sectioned, placed on glass slides, stained with hematoxylin and eosin (H&E), digitally scanned on an Aperio AT2 scanner (Leica Biosystems, Inc) creating whole slide images (WSIs), uploaded to the Histowiz cloud platform and evaluated by a senior board-certified research pathologist in a blinded fashion. The following measures were assessed: “Inflammation Score” (presence of inflammatory cell infiltrates and the approximate area containing macrophages, lymphocytes, plasma cells, and neutrophils), “Gland Loss Score” (proportion of mucosa with crypt epithelial and remaining gland epithelial loss), “Erosion/Ulceration Score” (loss of surface epithelium based on the estimated percent of the mucosa that was affected), “Mucosal Thickness Measurement” (measured in a non-tangential area of the section that best represents the overall mucosal thickness for that section), “Hyperplasia Score” (derived from the mucosal thickness measurement), “Histopathology Sum Score” (a sum of inflammation, gland loss, erosion, and hyperplasia scores), “Lymphoid Aggregate Count” (the number of definite mucosal lymphoid aggregates including GALT, Peyer’s patches, and defined inflammatory aggregates).

### Free-floating immunohistochemistry in brain and colon sections

For immunohistochemistry, free-floating brain sections were first washed with PBS, then permeabilized by incubation with 0.3% Triton X-100 in PBS while lightly shaking for 4 × 15 minutes at room temperature. Tissue was then blocked at room temperature for 2 hours in 0.3% Triton X-100, 10% Normal Donkey Serum (Jackson Immuno Research Laboratories # 017-000-121) in PBS, followed by overnight incubation (16-18 hours) at 4°C with primary antibodies diluted in 0.3% Triton X-100, 5% Normal Donkey Serum. The next day, sections were rinsed 4 × 15 minutes in PBS, then incubated for 120 minutes at room temperature in the appropriate secondary antibodies in a similar 5% serum cocktail as for primary antibodies. Lastly, sections were rinsed 4 × 10 min in PBS, incubated with DAPI (Sigma-Aldrich, # D8417), mounted in PBS and coverslipped with Fluoromount-G (Southern Biotechnology, # 100241-874), then left to cure for at least 24 h before imaging.

The following primary antibodies were used: guinea pig anti-c-Fos (1:1000, Synaptic Systems, # 226308, previously # 226004 but no difference was observed between product numbers), rabbit anti-VIP (1:1000, Immunostar, # 20077), rabbit anti-PV (1:2000, Swant, # PV27), rat anti-CD68 (1:1000, Bio-Rad, # MCA1957, clone FA-11), rabbit anti-Iba1 (1:500, Wako, # 019-19741), rabbit anti-SST (1:3000, BMA Biomedicals, # T4103), mouse anti-PV (1:2000, Swant, # PV235), mouse anti-gephyrin (1:500, Synaptic Systems, # 147011), rabbit anti-Homer1b/c (1:500, Synaptic Systems, # 160023), mouse anti-Synaptotagmin-2 (1:500, Zebrafish International Resource Center, # znp-1), rabbit anti-NeuN (1:1000, Millipore, # ABN78), guinea pig anti-VGluT2 (1:2000, Millipore, # AB2251), biotin-conjugated Wisteria floribunda agglutinin (1:2000, Sigma-Aldrich, # L1516), chicken anti-GFP (1:1000, Aves Labs, # GFP-1020), guinea pig anti-Brevican (1:2000, generously provided by Dr. Renato Frischknecht, Friedrich-Alexander-Universität), rabbit anti-cleaved Brevican (BCneoRT; 1:1000, generously provided by Dr. Renato Frischknecht, Friedrich-Alexander-Universität).

For CD45 immunohistochemistry in colon sections, free-floating tissue sections were first washed with PBS, then permeabilized by incubation with 0.3% Triton X-100 in PBS (PBST) while lightly shaking for 4 x 15 minutes at room temperature. Tissue was then blocked at room temperature using 3% bovine serum albumin (BSA; Sigma-Aldrich, # A9647) in PBST, followed by overnight incubation (16-18 hours) with anti-mouse CD45 Antibody pre-conjugated with Alexa Fluor® 647 (Biolegend, # 103123) diluted 1:100 with 3% BSA in PBST at 4°C. The next day, tissue was rinsed 4 x 10 minutes in PBS, incubated with DAPI, and subsequently rinsed in PBS before slide mounting in PBS. Coverslips were mounted with Fluoromount-G and allowed to cure for at least 24 h before imaging.

The following secondary antibodies were used (all at 1:500 dilution): donkey anti-rabbit Alexa Fluor® 546 (Thermo, # A-10040), donkey anti-guinea pig Alexa Fluor® 488 (Jackson Immuno, #706-545-148), goat anti-rat Alexa Fluor® 546 (Thermo, # A-11081), donkey anti-rabbit Alexa Fluor® 647 (Thermo, # A-31573), goat anti-mouse-IgG1 Alexa Fluor® 647 (Thermo, # A21240), Streptavidin Alexa Fluor® 647 (Thermo, # S-32357), goat anti-mouse-IgG2a Alexa Fluor® 488 (Thermo, # A-21131), donkey anti-rabbit Alexa Fluor® 488 (Thermo, # A-21206), goat anti-mouse-IgG1 DyLight™ 405 (BioLegend, # 409109), donkey anti-rabbit DyLight™ 405 (Jackson Immuno, # 711-475-152), goat anti-mouse-IgG1 Alexa Fluor® 546 (Thermo, # A-21123). Biotinylated WFA was applied together with the primary antibodies and revealed with streptavidin-Alexa Fluor 647. Within each experiment, sections from control and experimental animals were processed in the same batch with the same antibody cocktails, and both image acquisition and quantification were performed blind to treatment group.

### Confocal fluorescence microscopy

All confocal fluorescence microscopy was performed using a Zeiss LSM 900 laser scanning confocal microscope running Zeiss ZEN 3.4. Objectives used varied by application and included a Zeiss Plan-Apochromat 20x air objective (NA=0.8; CD45 and c-Fos analyses), a Zeiss Plan-Apochromat 40x oil immersion objective (NA=1.30; microglial, PV protein, WFA analyses), or a Zeiss Plan-Apochromat 63x oil immersion objective (NA=1.40; Brevican neo and synapse analyses). Within each experiment, laser lines and power, detector gain, pinhole, pixel size, z-step and averaging were held constant across all animals and groups, and sections from control and experimental mice were imaged in the same session.

### Quantification of CD45+ colon tissue

5-8 total sections of distal colon were analyzed per animal to account for within-animal variability in CD45 expression. Axial plane thickness was 1.4µm for all channels. A single image of CD45 and DAPI were acquired per section and saved as CZI files. Images were opened in FIJI/ImageJ and the entirety of the mucosal layer for each image was manually contoured using the DAPI channel. The CD45 channel was then thresholded manually (experimenter blind to experimental group) using the same threshold for all sections within an imaging batch and converted to a binary mask. The percent of mucosal area containing the CD45 mask was measured for each image. Within each animal, a ROUT outlier test (*75*) was applied to the per-section values at Q = 5%. The remaining section values were averaged to give one mean percentage of CD45+ mucosal area per animal. For comparison across cohorts, per-animal values are expressed as a percentage of the mean of the control group processed in the same batch.

### Quantification of c-Fos in interneuron populations

To quantify c-Fos+ VIP+ and SST+ interneurons, tiled images of the mid-posterior IC were acquired as single 1.4 µm optical sections. Using QuPath (version 0.5.0) (*76*), all VIP+ and SST+ cells were manually identified and quantified. For SST, c-Fos co-localization was performed in a semi-automated manner. First, regions of interest (ROI) were outlined using the “closed polygon annotation” function, in accordance with the Allen Coronal Mouse Brain Reference Atlas. Then, the “Positive Cell Detection” function was used to identify c-Fos+ cells. When using this function, the settings were first optimized to include all DAPI^+^ nuclei without segmentation that would break a single nucleus apart or group multiple nuclei together as one object. These objects would be called the “nucleus” for the cell detection. Then, a c-Fos+ threshold was optimized based on the mean AF488 intensity within the “nucleus” object. A single set of parameters was used across all ROIs on a single section. For VIP, c-Fos co-localization was performed entirely manually by an experimenter blinded to treatment. VIP/SST signal where a DAPI+ nucleus was not visible in-plane was excluded from analysis, because c-Fos is localized primarily to the nucleus, so including cells whose nucleus lay outside the optical section would artificially lower the estimated fraction of c-Fos+ interneurons. None of the cells lacking a visible DAPI+ nucleus showed c-Fos signal above background. For the VIP+ and SST+ analyses, the region of interest was the entire mid-posterior IC (layers 1-6; granular, dysgranular and agranular subregions), which maximized the number of SST+ cells sampled, as these are sparsely distributed across cortical layers and insular subregions. For VIP+ cells, 1-2 sections per animal were analyzed and 25-111 cells were sampled per animal. For SST+ cells, 1-2 sections per animal were analyzed and 31-168 cells were sampled per animal. Counting was performed blind to treatment group and animals with less than 25 total VIP+ or SST+ cells (i.e., c-Fos+ and c-Fos-combined) were excluded from analysis. No other exclusionary criteria were applied.

To quantify c-Fos+ PV+ interneurons, single 1.4 µm optical sections of layer 4 of the granular mid-posterior IC were acquired. Using Fiji/ImageJ, all PV+ cells were manually identified and quantified, as were cells where PV+ cell soma was colocalized with c-Fos+ signal. For the same reasons as above, PV signal where a DAPI+ nucleus was not visible in-plane was excluded from analysis, in order to avoid artificially lowering the estimated fraction of c-Fos+ interneurons. None of the cells lacking a visible DAPI+ nucleus showed c-Fos signal above background. Layer 4 of the granular mid-posterior IC was chosen because PV+ cell density is highest there. For PV+ neurons, 2-3 sections per animal were analyzed and 28-130 cells were sampled per animal. Counting was performed blind to treatment group. For the V1 analysis, c-Fos+ PV+ interneurons were quantified in layer 4 of the primary visual cortex (V1) by the same procedure.

### PV protein expression and perineuronal net analyses

PV protein expression and normalized perineuronal net (WFA) volume were quantified as described previously (*17*). Briefly, WFA and PV were visualized using AF647 and AF488, respectively, and confocal image stacks (40× oil immersion objective, NA 1.30, 0.33 µm step size) were reconstructed and analyzed with Imaris (Bitplane, version 10.1). PV+ neurons whose soma was fully contained within the section, in fields with optimal staining, were reconstructed in three dimensions. Three-dimensional isosurfaces were created around each PV-neuron soma using the ‘‘create surface’’ tool and soma volume and labeling intensities were quantified automatically. The threshold was set to include as much of the neuron as possible while excluding background, and was held constant across all animals within an experiment. A size filter was applied to exclude PV cells that were partially reconstructed or significantly out-of-stack. The PV+ surface was then dilated by 1 µm to define a perisomatic domain extending into the extracellular space. Within this domain, three-dimensional isosurfaces were constructed on the WFA channel, and WFA volume was normalized to the volume of the PV+ soma. Measurements were made in layer 4 of the granular mid-posterior IC, the same region used for c-Fos quantification, and 4-15 outliers were removed (ROUT test, Q=1%). Analysis of Brevican and BCneoRT staining around PV+ cells followed an identical protocol, using a 63x oil immersion objective (NA 1.40) to visualize PV, Brevican and BCneoRT using AF647, AF488 and AF546, respectively; the log2 Brevican/BCneoRT ratio was computed per cell from the two normalized volumes.

### Slice electrophysiology

Electrophysiology data were collected in three experiments, and all recordings were performed in the posterior IC: (i) Recordings of excitatory and inhibitory postsynaptic currents (miniature and spontaneous) in pyramidal and PV+ cells of WT mice after 2 or 4 days of 5% DSS. 2 weeks prior to recording, mice were injected with AAV2/9-S5E2-GFP-fGFP (200nL, 2x10^13^ GC/mL titer) bilaterally in mid-posterior IC. (ii) Recording of PV+ cells to measure intrinsic properties and synaptic currents in *TRAP2;Ai75* mice after 4 days of 3.5% DSS, with tamoxifen (150 mg/kg) administered on day 2. 2 weeks prior to recording, mice were injected with AAV2/9-S5E2-GFP-fGFP (200nL, 2x10^13^ GC/mL titer) bilaterally in the mid-posterior IC. (iii) Recording of pyramidal neurons to measure intrinsic properties in *TRAP2;Ai75* mice 5-6 days after the onset of 5% DSS, with tamoxifen (150 mg/kg, i.p.) administered on day 2. These mice did not receive AAV injection. The recordings of intrinsic properties were done in *TRAP2;Ai75* mice with a slightly higher tamoxifen dose to enrich for TRAPed neurons during the short labeling window, however, the low number of cells recovered did not allow comparison of TRAPed versus non-TRAPed neurons.

For all experiments, brain slices were prepared as previously described (*77*). In brief, mice were first anaesthetized with chloral hydrate (400 mg per kg, i.p.). After decapitation, the brains were removed rapidly and placed in ice-cold artificial CSF (aCSF) in which sucrose (252 mM) replaced NaCl equiosmotically (sucrose-ACSF) to limit excitotoxic swelling. 300 μm coronal slices were cut in sucrose-ACSF with an oscillating-blade vibratome (VT1200S, Leica). Slices were allowed to recover for around 1–2 h in sucrose-ACSF before commencement of recording. The slices were then placed into a submerged recording chamber in standard ACSF, and the bath temperature was raised to 32 °C. Slices were perfused with ACSF at 2-3 mL/min throughout recording. The standard ACSF (pH of around 7.35) was equilibrated with 95% O_2_/5% CO_2_ and contained 128 mM NaCl, 3 mM KCl, 2 mM CaCl_2_, 2 mM MgSO_4_, 24 mM NaHCO_3_, 1.25 mM NaH_2_PO_4_ and 10 mM d-glucose. Pyramidal cells in layer 4 and GFP+ PV+ cells in the granular mid-posterior IC were visualized under an Olympus BX51WI microscope using a ×60 infrared objective with infrared differential interference contrast videomicroscopy. An OLY-110 camera (ORCA-spark, Hamamatsu) was used to visualize neurons in the slice. Low-resistance patch pipettes (3–5 MΩ) were pulled from borosilicate glass (Warner Instrument) using a horizontal micropipette puller (P-1000, Sutter Instrument). Pipettes were filled with internal solution containing 115 mM K-gluconate, 5 mM KCl, 2 mM MgCl_2_, 2 mM Mg-ATP, 2 mM Na_2_ATP, 10 mM Na_2_-phosphocreatine, 0.4 mM Na_2_GTP and 10 mM HEPES, calibrated to pH 7.33. This K-gluconate-based internal solution was used for current-clamp measurements of intrinsic properties and for recording excitatory currents. For inhibitory postsynaptic current recordings, pipettes were filled with the solution contained (in mM): 125 Cs gluconate, 10 HEPES, 5 BAPTA-Cs, 10 sodium phosphocreatine, 2.38 CaCl2, 4 Mg-ATP and 0.3 Na2-GTP (pH 7.33). Excitatory and inhibitory postsynaptic currents were isolated by clamping neurons at −70 or 0 mV, respectively. Whole-cell patch-clamp recording was performed with an Axoclamp-2B amplifier (Axon Instruments), and the output signal was acquired using the pCLAMP 9.2 (Molecular Devices) /Digidata 1322A software. Series resistance, monitored throughout the experiment, was 4-8 MΩ. Cells were discarded if the series resistance rose above 8 MΩ. Liquid junction potential was not corrected. Miniature currents were recorded in the presence of tetrodotoxin (TTX,1 µM) bath-applied for at least 3 min before recording.

Intrinsic properties were measured rapidly after obtaining whole-cell configuration, by injecting a series of depolarizing and hyperpolarizing 500 ms current steps at 0.5 Hz. Resting membrane potentials were measured just after patched membranes were ruptured by suction. Input resistance was determined by passing hyperpolarizing current pulses inducing voltage shifts of 5-15 mV negative to resting membrane potential. Rheobase was determined by initially injecting positive current generating near-threshold 15-25 mV depolarizations and subsequently applying 1 pA current steps until APs were elicited. Rheobase was considered to have been reached once three such steps consecutively produced an AP, with the lowest value of the AP-generating injected currents used. AP amplitude was calculated as the difference in membrane potential between AP threshold and the peak. Maximum firing rate was determined from 20 pA current steps as the largest number of action potentials elicited within a single 500 ms step.

Synaptic currents were analyzed with Mini-Analysis (Synaptosoft, Decatur, GA, USA) and characterized by amplitude, rise time, duration and area under the curve (in fC). We used detection parameters of 5 pA for event amplitude threshold, area threshold of 50 fC, 10 ms for searching for a local maximum, 5 ms to search for baseline before peak, 50 ms to search for decay time and 0.37 as the fraction of peak amplitude to find a decay time. For each cell, three 20 s traces were recorded and the mean of the three (60 s in total) was used for analysis. Data were acquired and analyzed blind to the experimental design and expected outcomes, using detection parameters fixed in advance and applied identically to all cells.

### Quantification of synapses

We quantified the synapse density of Syt2+/Gephyrin+ and VGluT2+/Homer1b/c+ synapses onto cell somas (either PV+ or NeuN+) using a semi-automated analysis of single-plane images described previously (*78*) (citation includes representative FIJI/ImageJ scripts). Briefly, all channels were isolated and the “despeckle” and “smooth” functions in Fiji were applied. Next, a manual threshold was determined and a mask was created for each channel. From the PV or NeuN channel mask, the cell body plus the immediately adjacent 0.5 µm was selected and smoothed (using the “Enlarge” function to erode the outline by 2.5 µm and then dilate it by 3.0 µm) to generate a “soma” mask, whose perimeter was automatically measured. For bouton segmentation, the “watershed” function was applied to the presynaptic channel (Syt2 or VGluT2) and the postsynaptic channel (Gephyrin or Homer1b/c). We then used the “Analyze Particles” function (where the minimum size for presynaptic and postsynaptic structures was 0.20 µm^2^ and 0.10 µm^2^, respectively) to generate pre- and postsynaptic masks. The masks were merged, converted to 8-bit, and the overlap between presynaptic boutons, postsynaptic clusters and the soma mask was detected automatically with an automatic threshold and the “Analyze Particles” tool. To get final estimates of synapse density per cell, the number of soma/presynapse/postsynapse colocalized puncta per plane was normalized to the perimeter of the soma. Analyses were performed in layer 4 of the granular mid-posterior IC, with thresholds held constant across groups within an experiment, and blind to treatment group.

### Plasmid production, cell culture, transfection, and AAV production

The following AAVs were obtained directly from Addgene: AAV2/9-hSyn-DIO-hM3D(Gq)-mCherry (44361-AAV9), AAV2/9-hSyn-hM3D(Gq)-mCherry (50474-AAV9), AAV2/9-S5E2-GFP-fGFP (135631-AAV9), AAV2/9-hSyn-DIO-hM4D(Gi)-mCherry (44362-AAV9), AAV2/9-hSyn-DIO-mCherry (50459-AAV9). The plasmid pAAV-RAM-d2TTA::TRE-EGFP-WPREpA was obtained from Addgene (84469) and packaged in-house at Yale University. The plasmid pAAV-hDlx-PSAM^4^-5HT3 LC-IRES-mEGFP-WPRE3 was cloned in conjunction with VectorBuilder Inc. and packaged in-house at Yale University. The plasmid vectors pAAV-hDlx-K_ir_2.1-T2A-EGFP-WPRE3 and pAAV-hSyn-fDIO-Casp3-T2A-TEV-WPRE were cloned and packaged in conjunction with VectorBuilder Inc. Virus titers ranged from 10^9^ to 10^13^ GC/mL and are stated below for each AAV. The plasmids S5E2-GCaMP6f and BiPVe3-GCaMP6f were obtained from Gord Fishell and packaged in-house (7.2 × 10^9^ and 4.34 × 10^13^ GC/mL, respectively). Both enhancers were characterized in the IC and in primary somatosensory cortex (S1), and S5E2 was selected for all subsequent experiments because it labeled IC PV+ neurons with higher specificity.

For AAVs packaged in-house at Yale University, HEK293 cells (ATCC, # CRL-3216) were cultured in Dulbecco’s Modified Eagle’s medium (DMEM, Gibco #11965-092) supplemented with fetal bovine serum (Cytiva #SH30541.03, final concentration of 5%), Penicillin-Streptomycin (Gibco #15140122, final concentration of 100 U/mL), and GlutaMax (Gibco #35050061, final concentration of 1X). Cultures were maintained at 37 °C in a humidified incubator (PHCbi, # MCO-170AICUVHL-PA) containing 5% CO_2_. For AAV packaging and production, HEK293 cells were seeded on 15 cm plates and the plasmid of interest (6 µg/dish) was co-transfected with the following plasmids using polyethylenimine (300 µg/dish, Thermo # 43896-01): the AAV2/9 rep/cap plasmid (pAAV2/9n, Addgene # 122865; 6 µg/dish) and the adenoviral helper plasmid ([pAdDeltaF6, Addgene # 112867]; 18 µg/dish). Five days after transfection, cells and medium were harvested separately. Virus remaining in the medium was precipitated with polyethylene glycol 8000 (Promega # V3011), while the cell pellet was resuspended in freezing buffer (150 mM NaCl, 50 mM Tris, pH 8.0) and lysed by three freeze-thaw cycles; the two fractions were then combined and treated with Benzonase (Millipore, #70664-3 75 U/dish) and DNase 1 (Sigma-Aldrich, #D5307, 1 U/dish) at 37°C for 30 minutes. The supernatants were collected following centrifugation at 4000g for 10 minutes at room temperature (∼21°C), then layered onto discontinuous iodixanol gradients (15/25/40/60%; OptiPrep, Stemcell, # 07820) for ultracentrifugation (Type 70 Ti Beckman Rotor) at 58,400 r.p.m. for 2.25 h at 10 °C. The 40% iodixanol fraction containing the AAVs was collected and underwent ultrafiltration with DPBS supplemented with 1 mM MgCl2 and 2.5 mM KCl in an Amicon Ultra Centrifugal Filter (Millipore, #UFC910024). Ultrafiltration was performed six times at 4,000 r.p.m. for 30 min at 4 °C. Titers, reported throughout as genome copies (GC) per mL, were determined by qPCR against the AAV2 ITR sequence using the following primers: Fw-5’-GGA ACC CCT AGT GAT GGA GTT and Rv-5’-CGG CCT CAG TGA GCG A.

### Stereotaxic AAV injections

Mice were anesthetized with isoflurane (4% induction, 0.5-2% maintenance) in 100% O2 and held at a surgical plane, with body temperature maintained at 37 °C using a heated surgical platform. Sterile eye lubrication was applied. Mice received pre-operative analgesia (meloxicam, 5 mg/kg, i.p.; bupivacaine, 2 mg/kg, s.c.), and hair around the incision site was removed with depilatory cream before sterilization. Aseptic surgical technique was followed for all surgical procedures. A small incision (<1 cm) was made in the scalp, and measurements were taken to confirm a level head orientation. We targeted our injections in mid-posterior IC (*62*) using the following coordinates relative to Bregma: AP −0.10 mm, ML ±4.00 mm, DV −3.85 mm. Injections were bilateral or right-hemisphere only depending on the experiment, as specified for each experiment below and in the figure legends. To achieve this, we first drilled a small (<1 mm) craniectomy above each site. Because of the curvature of the skull at this lateral position, the DV coordinate was reached by inserting the glass microinjector tip to a depth of 2.40 mm ventral to the pial surface rather than measuring from Bregma. AAV injection was both preceded and followed by a 2-minute delay to account for disruption of parenchyma by insertion and to allow proper diffusion of AAV from the syringe tip after injection, respectively. All injections were performed at 2 nL/s; the total volume and viral titer for each construct are given below. Injections were made through a pulled glass micropipette driven by a Nanoject III Programmable Nanoliter Injector (Drummond Scientific). Injection sites were verified post hoc on the sections used for analysis: no animals had off-target or absent expression and therefore no animal was excluded. Upon completion of all injections, the skin was closed with 5-0 monofilament sutures and animals were kept on a heating pad until complete recovery from anesthesia. Post-operative analgesia consisted of daily meloxicam (5 mg/kg) for 3 days.

### Robust Activity Marking (RAM)

48 h before AAV injection, regular chow was replaced with ad libitum access to a doxycycline diet (Bio-Serv, # S3888). Through the surgical procedure outlined above, we injected AAV2/9-RAM-d2TTA::TRE-EGFP-WPREpA (150 nL, 7x10^12^ GC/mL titer) bilaterally in the mid-posterior IC. For the 5% experiment, an mCherry AAV was co-injected. Animals remained on a doxycycline diet during recovery. On the 10^th^ post-operative day, the doxycycline diet was replaced with a standard chow diet and drinking water was replaced with 5% or 3.5% DSS. Neither experiment involved tamoxifen. Brains and colons were perfusion-fixed and extracted as explained below after 4 days of DSS exposure. VIP-Cre heterozygous were used for the 5% experiment.

#### Quantification of RAM GFP+ neuron density

An Olympus VS200 slide scanner with 10x Olympus UPLXAPO objective (NA=0.4) and Hamamatsu ORCA-Fusion camera was used to acquire tiled widefield images of the mid-posterior IC at the site of AAV injection. Four serial coronal sections (1:8 serial interval) per animal were imaged, spanning the infected injection site (approximately +0.3mm (i.e., anterior) to -0.7mm (i.e., posterior) relative to Bregma). GFP+ cells within the injection site were manually quantified using QuPath (v0.7.0). For the 5% DSS experiment, regions of interest (ROIs) for quantification were drawn, constrained to the portions of insular cortex that expressed a co-injected constitutive mCherry-expressing AAV (AAV2/9-hSyn-hM3D(Gq)-mCherry). GFP+ neurons were then counted manually. GFP+ cell counts and ROI area were pooled across sections within each animal and density was calculated as the total number of GFP+ cells per mm² of ROI area for each animal. 1 control mouse was excluded due to abnormal expression of constitutive mCherry AAV (suggestive of double volume injection). For the 3.5% DSS experiment, tiled confocal images of the mid-posterior IC were acquired from five sections per animal, spanning +0.1mm (i.e., anterior) to -1.0mm (i.e., posterior) relative to Bregma to sample through the volume of the injection site. GFP+ neurons were counted manually in ImageJ within a region of interest comprising all layers and subregions of IC, and density is reported as GFP+ cells per mm² of section area. For both the 5% and 3.5% DSS experiments, ROI drawing and counting were performed blind to treatment group.

### LPS injection

Mice received 2 i.p. injections of LPS (1 mg/kg; from Escherichia coli O111:B4; Sigma-Aldrich, # L2630) or phosphate-buffered saline (PBS) once every 24 h, and brains were collected 12 h after the final injection. LPS was prepared as a 5 mg/mL stock solution and diluted to a working concentration of 0.1 mg/mL in PBS before administration.

### Microglia analyses

#### Quantification of microglia density

Microglial density was quantified in QuPath (version 0.5.0). The IC was outlined as the region of interest and its area measured; candidate cells within the region were detected with the QuPath cell-detection command, and Iba1+ microglia were then labeled and counted manually in layers 1-6. Density is reported as Iba1+ cells per mm^2^ of section area.

#### Quantification of lysosome occupancy and PV puncta in microglia

For lysosome occupancy in microglia, confocal stacks (40× oil immersion objective, NA 1.30, 0.38 µm step size, 7-15 μm total depth) were analyzed with Imaris version 10.1.0 or 10.1.1. Three-dimensional isosurfaces were created based on the confocal stack image of Iba1+ microglia using the Surface tool. CD68+ lysosomes were reconstructed in the same way, and microglial volume and the fraction of that volume occupied by CD68+ surfaces were quantified automatically. The quantification of engulfed PV+ puncta was performed as described before (*78*). Briefly, Syt2+ presynaptic puncta were segmented as spots using the Spots tool in Imaris. The diameter threshold was set to 0.8 μm as described before (*78*). Spots whose distance to the microglial surface was less than 0 μm, that is spots lying inside the surface, were classified as “engulfed”. These spots were assigned to individual microglia using the “Spots Split Into Surface Objects” XTension in Imaris, and the number of engulfed spots per microglial cell was recorded automatically.

#### Sholl analysis of Microglia

Microglial arbors were reconstructed with the Filaments tool in Imaris. The soma reconstruction was adjusted manually to match its shape and to define the soma center point, and filaments were then traced using the machine-learning-assisted mode. The same trained classifier was applied to control and DSS images. Only fully contained reconstructions were retained, and branches that were not continuous in the original image were removed. Sholl analysis used a shell spacing of 3 μm, and the resulting intersection profiles were analyzed and plotted in R with a linear mixed-effects model (random intercept per cell nested within animal), as recommended for Sholl data (*79*).

### Microglia depletion via dietary PLX5622 administration

Microglial depletion was achieved with the CSF1R inhibitor PLX5622 (MedChemExpress, # HY-114153), as previously reported (*78*). Mice were given ad libitum access to PLX5622-containing chow (1,200 mg PLX5622 per kg chow, i.e., 1,200 ppm, formulated in an AIN-76A base diet; Research Diets, # an-0002) for 11 days before the initiation of DSS. Both PLX5622-containing and control chow were stored at −20 °C protected from light, replaced in the cage every 2 days, and thawed 12 h before administration to limit degradation of PLX5622 by ambient light and room temperature. Animals remained on their assigned diet until tissue collection, after 4 days of 5% DSS or control water.

### Subdiaphragmatic vagotomy (SDV)

A single dose of extended-release buprenorphine (Ethiqa XR, 3.25 mg/kg, s.c. # 072117 Butler Animal Health Holding, Covetrus), followed by a local block (bupivacaine, 2 mg/kg s.c.; # AAJ6283506), was administered pre-operatively. Bilateral SDV was performed under continuous inhalation anesthesia with 2%-3.5% isoflurane using an inhalation small animal anesthesia apparatus on a sterile heated surgical surface, following a previously reported protocol (*80*). Briefly, the surgical site was disinfected, an incision of approximately 1 cm was made parallel to the costal arch, and the underlying liver tissue was carefully pushed upward using a small sterilized cotton ball moistened with sterile saline (0.9%). The dorsal and ventral branches of the vagus nerve, which run along the esophagus under the diaphragm, were carefully separated and severed. The liver tissue was returned to its original position, and sterile saline solution was injected into the abdominal cavity prior to layer-by-layer closure of muscle and skin layers using sterile monofilament surgical suture. Upon removal of anesthesia, thermal support was maintained until the animal could maintain unassisted sternal recumbency. For the sham operation, an identical surgical procedure was performed, but the dorsal and ventral branches of the subdiaphragmatic vagus nerve were not severed once exposed. All other aspects of the procedure were identical. Completeness of vagotomy was confirmed postmortem by inspection of the subdiaphragmatic esophagus and by gastric distension, and all animals had evidence of complete transection of both branches. SDV and sham mice were operated on and processed in parallel, received softened chow and were weighed daily. On the 7^th^ post-operative day, we administered either 5% DSS or H_2_O. Brains and colons were perfusion-fixed and extracted as explained above after 2 days of DSS or control H_2_O exposure.

### Trpv1+ sensory neuron ablation

*TRPV1-DTR* mice (*42*) were provided by I. Chiu (Harvard Medical School) and originated from M. Hoon (NIH). Genotyping was performed by PCR using genomic DNA isolated from mouse ear biopsies. The *DTR* allele was identified using generic EGFP primers obtained from the Mutant Mouse Resource and Research Center (MMRRC) at the University of California, Davis (UC Davis; 530-754-MMRRC). To ablate TRPV1-expressing neurons, Trpv1-DTR mice received daily i.p. injections of 200 ng diphtheria toxin (DT; Sigma-Aldrich, # D0664) in 100 μL PBS or 100 μL PBS alone (vehicle), for 21 consecutive days. 3-7 days after the last DT injection, mice were sensitized and challenged with HDM; brains and DRG were collected 48 h after sensitization and 48 h after challenge, as indicated in the relevant figures. In a separate cohort, DT- or vehicle-treated Trpv1-DTR mice received 5% DSS or water, and brains and DRG were collected 48 h after the onset of DSS. For immunostaining, mice were transcardially perfused with PBS followed by 4% PFA in PBS. Thoracic dorsal root ganglia (DRG, T1-T13) were dissected and postfixed for 2 hours in 4% PFA/PBS at 4 °C, incubated overnight at 4 °C with 30% sucrose in PBS, embedded in optimal cutting temperature compound (OCT, Tissue-Tek, PA), and stored at −80 °C. DRGs were then sectioned on a cryostat (Leica). Cryosections of 12 μm were cut, mounted on superfrost slide glass, and immunostained with the following antibodies: guinea pig anti-TRPV1 (1:1,000; Millipore, # AB5566) and donkey anti-guinea pig Alexa Fluor 488 (1:500; Jackson ImmunoResearch, # 706-545-148). Sections were mounted with Fluoromount-G and imaged with an LSM 900 confocal microscope. To measure heat sensitivity, mice were placed on a hot plate set at 55 °C, and the latency to hindpaw lifting, licking, or flinching was recorded, with a 60 s cutoff to prevent tissue injury. Ablation was validated by hot plate latency and by TRPV1 immunostaining in thoracic dorsal root ganglia.

### HDM model of respiratory allergy

We administered house dust mite (HDM; Dermatophagoides pteronyssinus, Stallergenes Greer, Item # XPB82D3A2.5, lot #s 413182, 434903) by intratracheal administration under anesthesia (*81*). HDM was resuspended in sterile 1x PBS at 4 mg/mL and stored at -80°C until use.

Immediately before use, the 4 mg/mL HDM stock was diluted 1:1 with sterile 1x PBS. Animals were anesthetized with i.p. injection of ketamine (100 mg/kg) and xylazine (10 mg/kg) and received 100 µg HDM in 50 µL. Animals were monitored and provided thermal support until cessation of anesthesia. Control animals were administered sterile 1x PBS without HDM. For “sensitization” experiments, only one dose of HDM or PBS was administered. For “challenge” experiments, a second dose of either HDM or PBS was administered exactly 7 days after sensitization. Brains were collected 2, 4 or 6 days after sensitization or after challenge, as indicated in each figure.

### Bronchoalveolar lavage and flow cytometry

Bronchoalveolar lavage (BAL) fluid was collected 2 days after HDM sensitization or 3 days after challenge by four sequential washes with 0.8 mL PBS (3.2 mL total instilled volume per mouse). BAL cells were centrifuged at 450 × g for 5 min and incubated with 1 mL ammonium-chloride-potassium (ACK) lysis buffer (Gibco, A10492-01) for 5 min to lyse erythrocytes. Lysis was quenched with 1 mL FACS buffer (PBS supplemented with 2% fetal bovine serum), and the cells were centrifuged and washed with 1 mL FACS buffer. To block nonspecific Fc receptor binding, cells were incubated in 10 µL FACS buffer containing purified rat anti-mouse CD16/CD32 antibody (Mouse BD Fc Block™, clone 2.4G2, 1:200, # 553142, BD Biosciences) for 5 min at room temperature. Cells were stained with a fluorophore-conjugated antibody cocktail containing PerCP-Cy5.5 anti-mouse CD4 (clone GK1.5, 1:200, #569847; BD Biosciences), BV605 anti-mouse CD8a (clone 53-6.7, 1:400, #563152; BD Biosciences), BV510 anti-mouse/human CD11b (clone M1/70, 1:200, #101263; BioLegend), PE/Cy7 anti-mouse CD11c (clone N418, 1:200, #117318; BioLegend), Alexa Fluor 488 anti-mouse TCRβ (clone H57-597, 1:200, #109215; BioLegend), BV711 anti-mouse Ly6G (clone 1A8, 1:200, #127643; BioLegend), BB515 anti-mouse Siglec-F (clone E50-2440, 1:200, #564514; BD Biosciences), and R718 anti-mouse TER-119 (clone TER-119, 1:200, #567363; BD Biosciences).Antibody staining was performed for 25 min at 4 °C, followed by two washes with 1 mL FACS buffer. Cells were resuspended in 200 µL FACS buffer and filtered through a 70-µm nylon mesh. Immediately before acquisition, DAPI was added at a final concentration of 0.2 ug/mL to identify nonviable cells. The entire sample volume was acquired on a Cytek Aurora spectral flow cytometer (Cytek Biosciences). Total cell numbers for each population were determined from the total number of events within the corresponding gate after exclusion of debris, doublets, and nonviable cells. Single-stained and unstained controls were used for spectral unmixing. Data were analyzed using FlowJo v10 and GraphPad Prism (version 9.0).

### Reactivation of neuronal engrams using *TRAP2* mice

We used the *TRAP2* mouse line to reactivate neurons that respond to peripheral immune challenge, as previously published (*4*, *46*). Briefly, we injected the mid-posterior IC of *TRAP2* mice with AAV2/9-hSyn-DIO-hM3D(Gq)-mCherry (2x10^12^ GC/mL, 150 nL), adhering to the stereotaxic surgical protocol outlined above. After 10-14 days of post-operative recovery, we administered 3.5% DSS for 4 days. Tamoxifen was dissolved in corn oil (20 mg/ml) at 37°C with sonication and stored at -80°C. Before injection, a 10 mg/ml dilution in corn oil was made. Tamoxifen was injected i.p. 48 h after the onset of DSS at 100 mg/kg body weight (except for some of the electrophysiology cohorts, where 150 mg/kg was used, see below). 4 weeks after the removal of DSS, we administered clozapine N-oxide (CNO; Tocris Bioscience, # 4936). CNO was resuspended in DMSO at 1 mg/ml, diluted in 0.9% saline to make a 0.1 mg/ml working solution, and injected i.p. every 12 h for 72 h at 1 mg/kg of animal weight. Tissue was fixed and extracted as described above 84 h after the first injection, that is 12 h after the last. Of note, repeated attempts to capture the immune engram using *TRAP2* mice crossed with *Ai75D* (Jackson # 025106) or *CAG-Sun1/sfGFP* (Jackson # 021039) were unsuccessful: variability among control animals was too high to permit reliable quantification.

### Validation of hDlx-selective enhancer AAVs

To measure the sensitivity and specificity of the hDlx enhancer in IC, mice were perfused 2 weeks after AAV injection and sections from the IC were stained as described above. Imaging was performed on a confocal microscope using a 20x objective and colocalization was quantified. Sensitivity was calculated as the percentage of GABA+ cells that were GFP+, and specificity as the percentage of GFP+ cells that were GABA+, counted manually in ImageJ.

### Interneuron manipulations during the encoding window

#### Pharmacological enhancement of GABAergic tone using Diazepam

TRAP2 mice were injected bilaterally in the mid-posterior IC with AAV2/9-hSyn-DIO-hM3D(Gq)-mCherry (2x10^12^ GC/mL, 150 nL per side), as described above. To pharmacologically mimic increased GABAergic inhibitory tone observed around day four of DSS, we administered diazepam (20 mg/kg in saline; Dash Pharmaceuticals, NDC # 69339-0137-05) i.p., following previously published dosing (*82*). Diazepam was administered every 12 h, beginning 36 h after the initiation of 3.5% DSS and ending 48 h after DSS withdrawal. During this time, tamoxifen was administered as described above (i.e., 48 hours after initiation of DSS).

#### Silencing of IC inhibitory interneurons with K_ir_2.1

We silenced inhibitory interneuron populations while utilizing our TRAP/Gq-based engram reactivation approach (described above) by co-injecting AAV2/9-hDlx-K_ir_2.1-T2A-EGFP-WPRE3 unilaterally into the right mid-posterior IC of TRAP2 mice, together with the AAV2/9-hSyn-DIO-hM3D(Gq)-mCherry described above (Kir2.1 virus 8x10^12^ GC/mL, 150nL). Control mice received the Gq virus alone, and the total viral load was matched between groups. After 11 days to allow expression, we administered 3.5% DSS. Tamoxifen was administered as described above (i.e., 48 hours after initiation of DSS).

#### Temporally selective activation of IC inhibitory neurons with PSAM^4^-5HT3 LC

We activated interneuron populations orthogonally to our TRAP2/Gq-based engram reactivation approach (described above) by co-injecting AAV2/9-hDlx-PSAM^4^-5HT3 LC-IRES-mEGFP-WPRE3 (6x10^12^ GC/mL, 150nL) unilaterally into the right mid-posterior IC of TRAP2 mice, together with AAV2/9-hSyn-DIO-hM3D(Gq)-mCherry as described above. Three weeks after surgery, we administered 3.5% DSS. Tamoxifen was administered as described above (i.e., 48 hours after initiation of DSS). To activate the PSAM4-5HT3 LC construct and drive interneuron activity, Varenicline (5 µg/mL; Tocris Bioscience, # 3754) was added to the DSS solution 24 h after the start of DSS and, once DSS was withdrawn, to the drinking water. Although the manipulation targets the encoding window (days 1-4), varenicline was maintained until 72 h after DSS removal to prevent encoding during the late inflammatory phase.

#### Targeted ablation of IC PV cells

We selectively ablated PV interneurons in the right mid-posterior IC while utilizing our Gq-based reactivation approach by co-injecting AAV2/9-hSyn-fDIO-Casp3-T2A-TEV-WPRE (6x10^12^ GC/mL, 150nL) unilaterally into the right mid-posterior IC of *TRAP2;PV-FlpO* mice, together with AAV2/9-hSyn-DIO-hM3D(Gq)-mCherry as described above. 14 days were allowed for ablation of PV+ neurons before 3.5% DSS was administered. Tamoxifen was administered as described above (i.e., 48 hours after initiation of DSS).

#### Silencing of VIP+ interneurons during colitis

We silenced VIP+ interneuron activity with time and region specificity by injecting AAV2/9-hSyn-DIO-hM4D(Gi)-mCherry (2x10^13^ GC/mL, 150nL) bilaterally into mid-posterior IC of *VIP-Cre* mice. Control animals received AAV2/9-hSyn-DIO-mCherry (2x10^13^ GC/mL, 150nL). After 11-12 days of recovery, we administered 5% DSS. To silence inflammation-induced VIP+ interneuron activity, we administered CNO (1 mg/kg, i.p.) every 12 h from 24 h to 84 h after the onset of DSS; control animals received the same CNO regimen. Tissue fixation and extraction (as described above) occurred 3 hours after the final CNO administration (i.e., on the fourth day of DSS administration).

### Statistical testing

All of the statistical details of experiments can be found in the figure legends, which include the statistical tests used, exact value of n, exclusion of any data, and what n represents. Data analysis was performed in GraphPad Prism (version 11.1.0) except where explicitly indicated. Data sets were first examined for normality with a Shapiro-Wilk test. If data were ordinal (e.g., feces pathology score) or non-normal (Shapiro-Wilk p < 0.050), groups were compared using a Mann-Whitney U test. If data were normally distributed, homogeneity of variance was checked using an F-test to compare group variances. If the F-test p<0.050, a Welch’s t-test was used to compare group means. Otherwise, a Student’s unpaired t-test was used to compare group means. Where the comparison concerned the distribution of a cell-level measurement rather than a group mean, such as perineuronal net volume or synapse density, groups were compared with a two-sample Kolmogorov-Smirnov test.

For comparisons among three or more groups, normality and homogeneity of variance were assessed as above. Normally distributed data were compared using a one-way ANOVA, followed by Tukey’s multiple comparisons test when all pairwise comparisons were of interest, or by Dunnett’s multiple comparisons test when each group was compared against a single control. Ordinal or non-normal data were compared using a Kruskal-Wallis test followed by Dunn’s multiple comparisons test. Experiments with two factors, such as microglial depletion combined with DSS treatment, were analyzed using a two-way ANOVA followed by uncorrected Fisher’s LSD for post hoc comparisons. Ordinal measures collected repeatedly across an experimental timeline, such as feces pathology scores, were compared between groups at each timepoint using multiple Mann-Whitney tests, and repeated measurements of the same group across timepoints were compared using a repeated-measures ANOVA.

Some datasets with nested or repeated observations were analyzed using linear mixed-effects models. Firing rate as a function of injected current was modeled with random intercepts and slopes per cell. Functional connectivity across the DSS time course was modeled with a group-by-time interaction term. For the functional ultrasound imaging dataset, connectivity values were tested against zero using one-sample t-tests, and within-group changes from baseline were tested using paired t-tests on the Fisher z-transformed change. Linear fits of pathology score and colon length against DSS duration were tested for a non-zero slope by linear regression. All tests were two-tailed, and p < 0.05 was treated as significant; exact p values are given in the figure legends where 0.05 < p < 0.10, and throughout for the functional ultrasound imaging analyses.

Animals were assigned to treatment groups at random, and control and experimental animals were processed, imaged and analyzed in parallel. Because DSS is administered in the drinking water, treatment was necessarily assigned at the level of the cage rather than the individual animal. However, treatment groups generally comprised mice from at least two cages, so that cage was not fully confounded with treatment, and control and DSS cages were housed on the same rack and were handled, perfused and processed in parallel. Image acquisition and quantification were performed blind to treatment group. Group sizes were chosen to match those used in previous studies with the same assays. All n values, the unit of replication and any exclusions are reported in the figure legends.

---

**Fig. S1.**
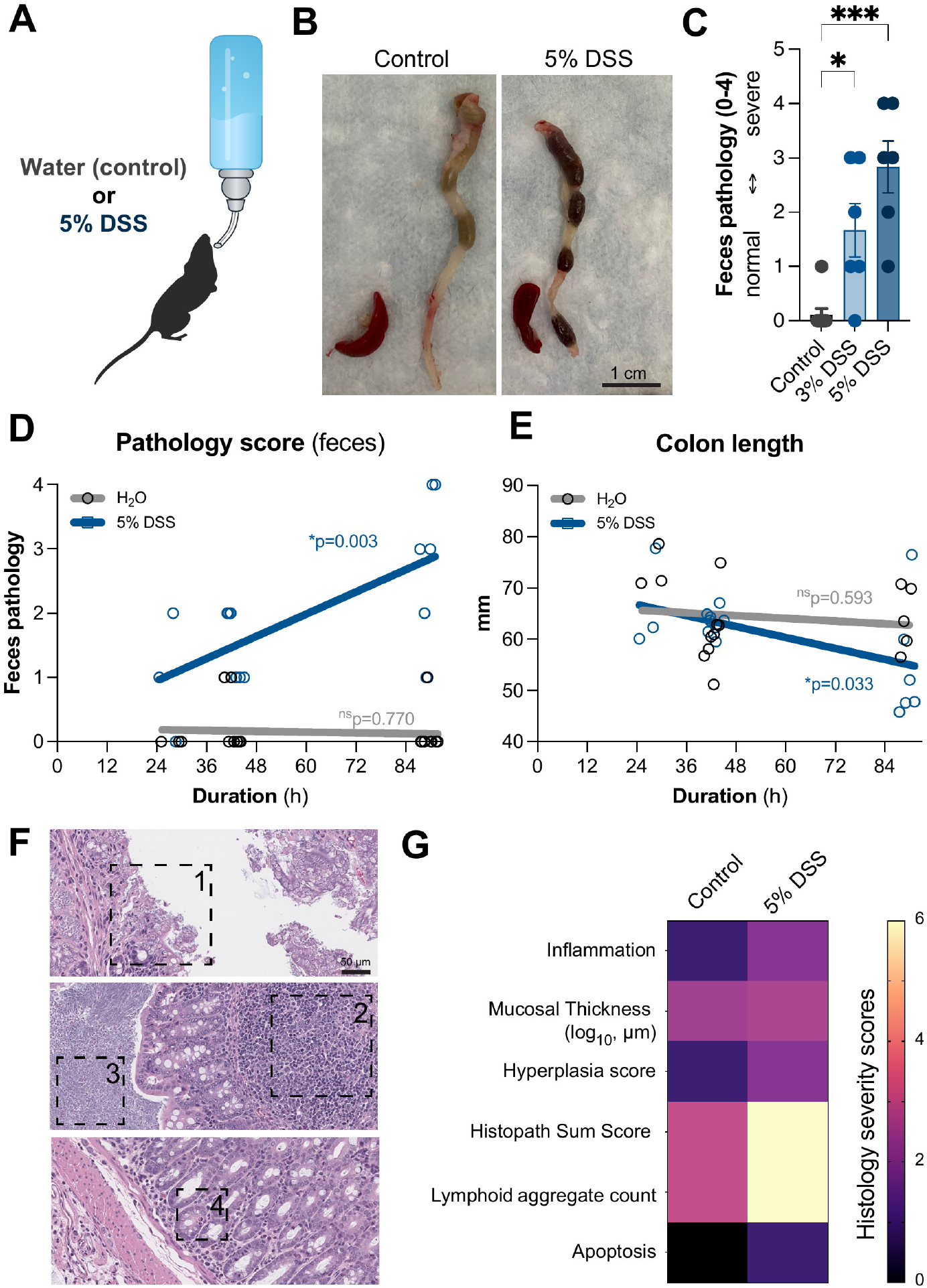
DSS administration induces dose- and time-dependent colitis. **(A)** Schematic of the DSS administration paradigm. **(B)** Representative colons from control and 5% DSS mice. **(C)** Feces pathology score (0-4, from normal to severe) in control mice and after 4 days of 3% DSS and 5% DSS (n = 6-9 mice/group). **(D-E)** Feces pathology score (D) and colon length (E) as a function of DSS duration (n = 36-38 mice/group, 3-6 mice/group/timepoint). Lines are linear fits; p values are for a non-zero slope. **(F-G)** Representative hematoxylin and eosin images of colon sections from a DSS mouse (F). Top panel: focal ulceration, epithelial erosion (1). Mid-panel: lymphoid aggregate, with lymphocytes, plasma cells (2); filamentous bacterial colonies (3). Bottom panel: Area with apoptosis (4). Histological severity scores (G) in a control and a 5% DSS mouse. *p<0.05, ***p<0.001; Kruskal-Wallis test with Dunn’s multiple comparisons test (C) and linear regression (D, E). Data are mean ± SEM.

**Fig. S2.**
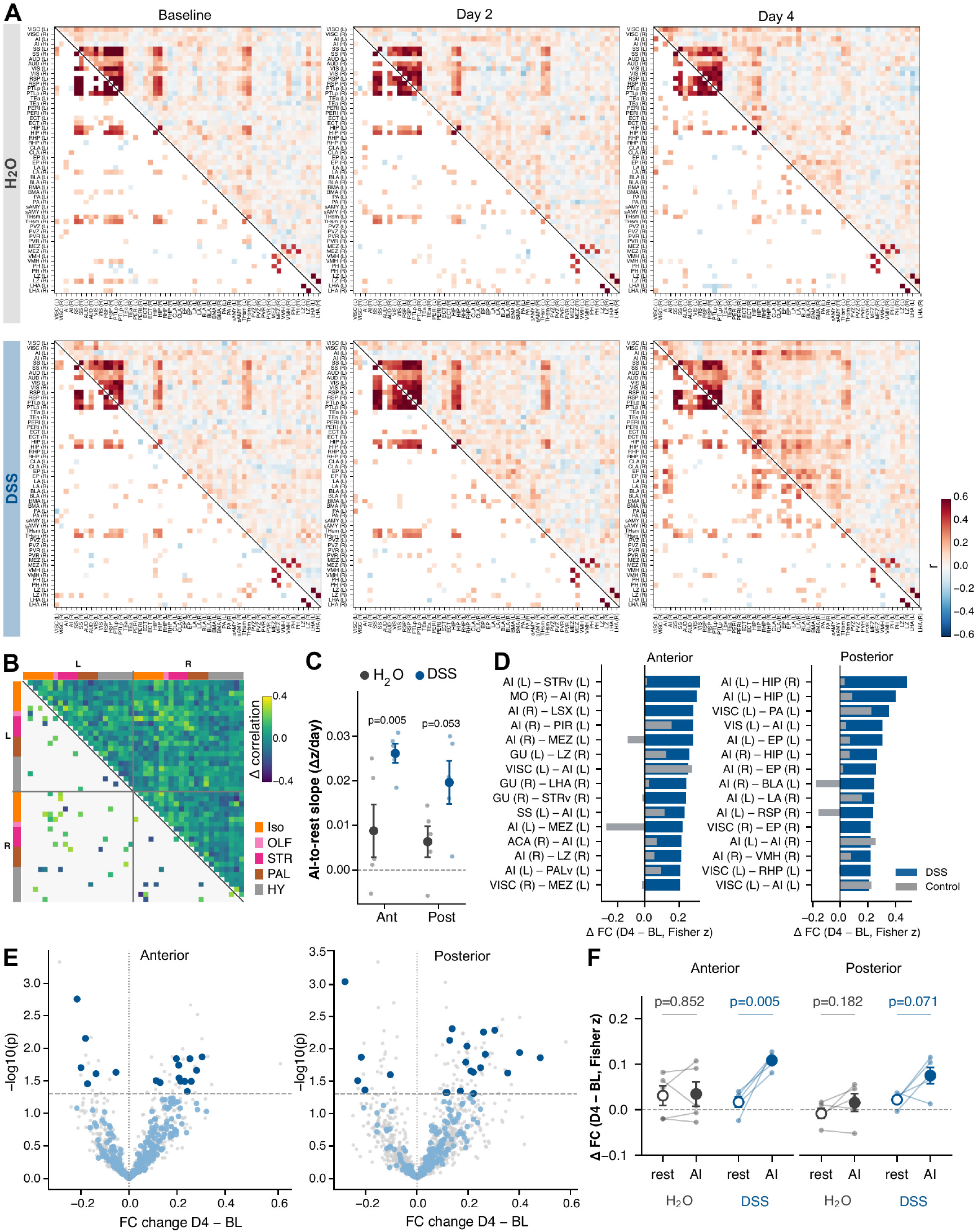
Functional connectivity reorganizes across the brain during colitis. **(A)** Group-averaged functional connectivity (Pearson r) in H_2_O and DSS mice at baseline, Day 2 and Day 4 (n = 5 per group; posterior plane). Each cell is the mean correlation between two regions. In each panel, the upper triangle shows all region pairs and the lower triangle shows only pairs significantly different from zero across that group’s mice (one-sample t-test, p < 0.05). **(B)** Matrices of the change in pairwise functional connectivity between anatomically defined regions of interest (ROIs), as in Fig. 1C, but for the anterior plane. Each cell is the change in the group-mean Pearson correlation between two ROIs (Δr). The upper triangle shows all region pairs; the lower triangle shows only pairs that changed significantly in DSS (within-group paired t-test on the Fisher z change, p < 0.05). Regions are ordered by left (L) and right (R) hemisphere and grouped within each hemisphere by Allen brain macroregion (see abbreviations below). **(C)** Rate of change of core insula-to-rest connectivity across the DSS time course (rate of recruitment), in the anterior and posterior planes. For each mouse, a line was fit to its core insula-to-rest connectivity (the mean Fisher z-transformed correlation of the left and right agranular insular cortex with all other regions) across baseline, Day 2 and Day 4, and the slope was taken (change in Fisher z per day). Faint dots are individual mice; the filled point and error bar are the group mean ± SEM (n = 5 per group). Linear mixed model, group by time interaction, two-tailed: anterior p = 0.005, posterior p = 0.053. **(D)** Top fifteen insula-involving region pairs with the largest increase in connectivity from baseline to Day 4 in DSS mice, for the anterior and posterior planes. Bars show the mean within-group change in Fisher z-transformed connectivity (Δ, Day 4 minus baseline) for DSS (blue) and H_2_O (gray) mice (n = 5 per group). Edges are labeled by their two regions using Allen atlas abbreviations. **(E)** Edge-level volcano plots of the within-DSS change from baseline to Day 4, for the anterior and posterior planes. For every region pair, the x-axis is the mean change in Fisher z-transformed connectivity across DSS mice, and the y-axis is the negative log10 of the p-value from a within-DSS paired t-test (Day 4 versus baseline). Insula-involving edges are highlighted (dark blue, p < 0.05; light blue, not significant); non-insula edges are gray. The dashed line marks p = 0.05. **(F)** Change in functional connectivity from baseline to Day 4 (Fisher z) between the core insula (AI) and all other regions, and among all region pairs that do not involve the insula (rest), for the anterior and posterior planes. Each line is one mouse; large symbols are group means ± SEM (n = 5 mice per group). p values test whether AI increased more than rest in the same mouse (two-tailed one-sample t-test on the per-mouse difference). **Region abbreviations.** ACA, anterior cingulate area; AHN, anterior hypothalamic nucleus; AI, agranular insular area; AUD, auditory areas; BLA, basolateral amygdalar nucleus; BMA, basomedial amygdalar nucleus; CLA, claustrum; ECT, ectorhinal area; EP, endopiriform nucleus; GU, gustatory areas; HIP, hippocampal region; LA, lateral amygdalar nucleus; LHA, lateral hypothalamic area; LPO, lateral preoptic area; LSX, lateral septal complex; LZ, hypothalamic lateral zone; MEZ, hypothalamic medial zone; MO, somatomotor areas; PA, posterior amygdalar nucleus; PALc, pallidum caudal region; PALd, pallidum dorsal region; PALm, pallidum medial region; PALv, pallidum ventral region; PERI, perirhinal area; PH, posterior hypothalamic nucleus; PIR, piriform area; PTLp, posterior parietal association areas; PVR, periventricular region; PVZ, periventricular zone; RHP, retrohippocampal region; RSP, retrosplenial area; SS, somatosensory areas; STRd, striatum dorsal region; STRv, striatum ventral region; TEa, temporal association areas; THsm, thalamus sensory-motor cortex related; VIS, visual areas; VISC, visceral area; VMH, ventromedial hypothalamic nucleus; sAMY, striatum-like amygdalar nuclei. Macroregions: Iso, isocortex; OLF, olfactory areas; CTXsp, cortical subplate; HPF, hippocampal formation; STR, striatum; PAL, pallidum; TH, thalamus; HY, hypothalamus. L and R denote left and right hemispheres.

**Fig. S3.**
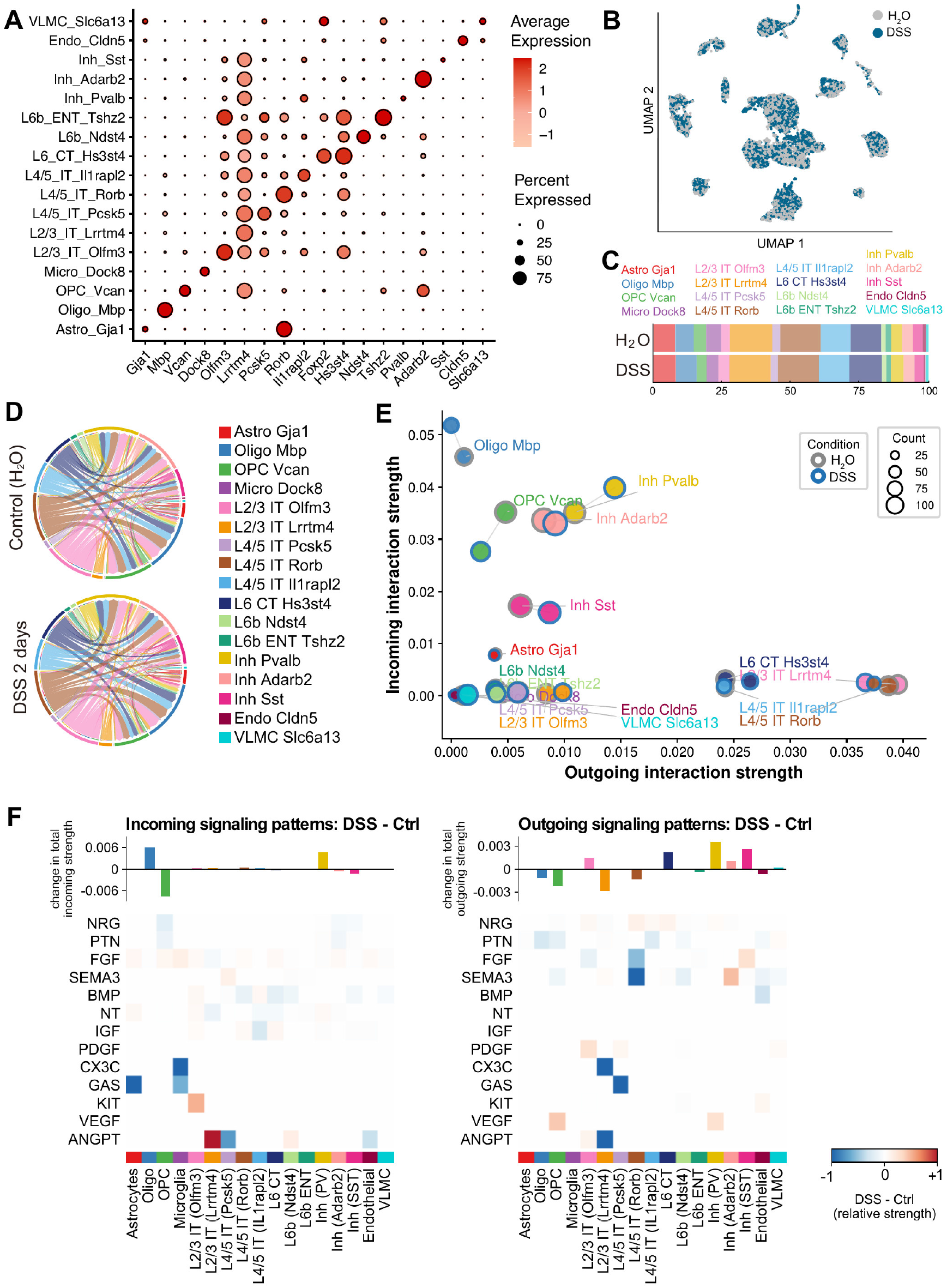
Cell-type identity, composition, and cell-cell communication in the IC following DSS colitis. **(A)** Dot plot of marker gene expression across IC cell types. Dot color indicates average scaled expression and dot size the percentage of nuclei expressing each gene. **(B)** UMAP of the integrated snRNA-seq dataset, split by condition. **(C)** Proportion of nuclei assigned to each cell type in control and DSS mice. **(D)** Chord diagrams of inferred cell-cell communication in control and 2-day DSS mice. **(E)** Incoming versus outgoing interaction strength for each IC cell type in control and DSS mice, inferred by cell-cell communication analysis. Point size indicates the number of inferred interactions. **(F)** Change in incoming (left) and outgoing (right) signaling strength between DSS and control for each signaling pathway and IC cell type. Bar plots above show the change in total interaction strength per cell type.

**Fig. S4.**
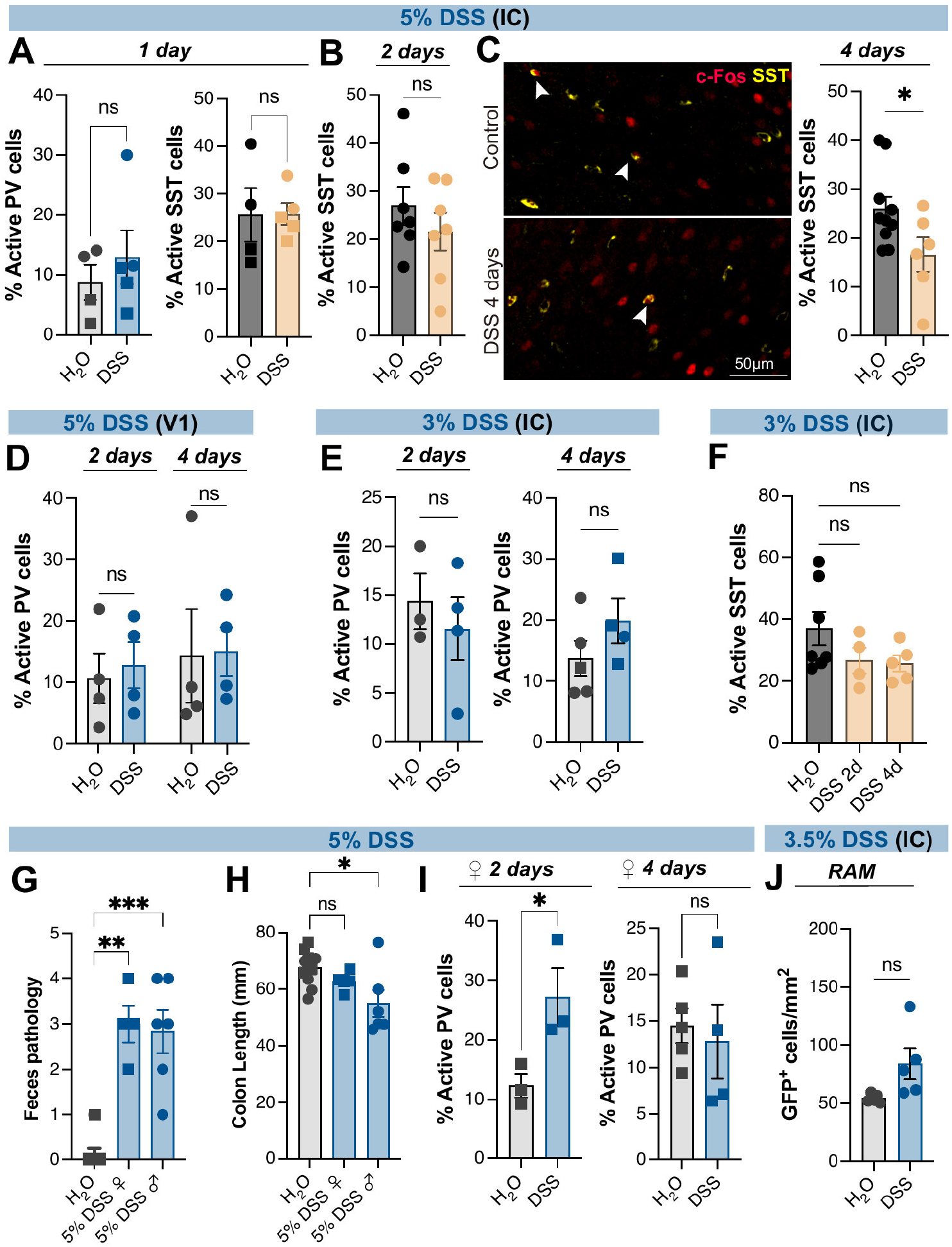
IC neuronal responses across time points, DSS doses and sexes. **(A)** Fraction of c-Fos+ PV+ and SST+ interneurons in the IC at 1 day of 5% DSS (n = 4-5 mice/group). **(B)** Fraction of c-Fos+ SST+ interneurons in the IC at 2 days of 5% DSS (n = 7 mice/group). **(C)** Representative images and fraction of c-Fos+ SST+ interneurons in the IC at 4 days of 5% DSS (n = 6-10 mice/group). Arrowheads indicate c-Fos+ SST+ interneurons. **(D)** Fraction of c-Fos+ PV+ interneurons in the primary visual cortex (V1) at 2 and 4 days of 5% DSS (n = 4 mice/group). **(E)** Fraction of c-Fos+ PV+ interneurons in the IC at 2 and 4 days of 3% DSS (n = 4-5 mice/group). **(F)** Fraction of c-Fos+ SST+ interneurons in the IC at 2 and 4 days of 3% DSS (n = 4-7 mice/group). **(G-H)** Feces pathology score (G) and colon length (H) in control and 5% DSS mice (n = 4-10 in G and 4-14 in H mice/group). **(I**) Fraction of c-Fos+ PV+ interneurons in the IC of female mice at 2 and 4 days of 5% DSS (n = 3-5 mice/group). **(J)** Density of RAM GFP+ neurons in the IC of control and 4-day 3.5% DSS mice, without tamoxifen (n = 5 mice/group). *p<0.05, **p<0.01, ***p<0.001; unpaired t-test (A-E, I), one-way ANOVA with Dunnett’s multiple comparisons test (F, H), Kruskal-Wallis test with Dunn’s multiple comparisons test (G), Welch’s t-test (J). Data are mean ± SEM; in this figure, circles and square symbols indicate males and females, respectively.

**Fig. S5.**
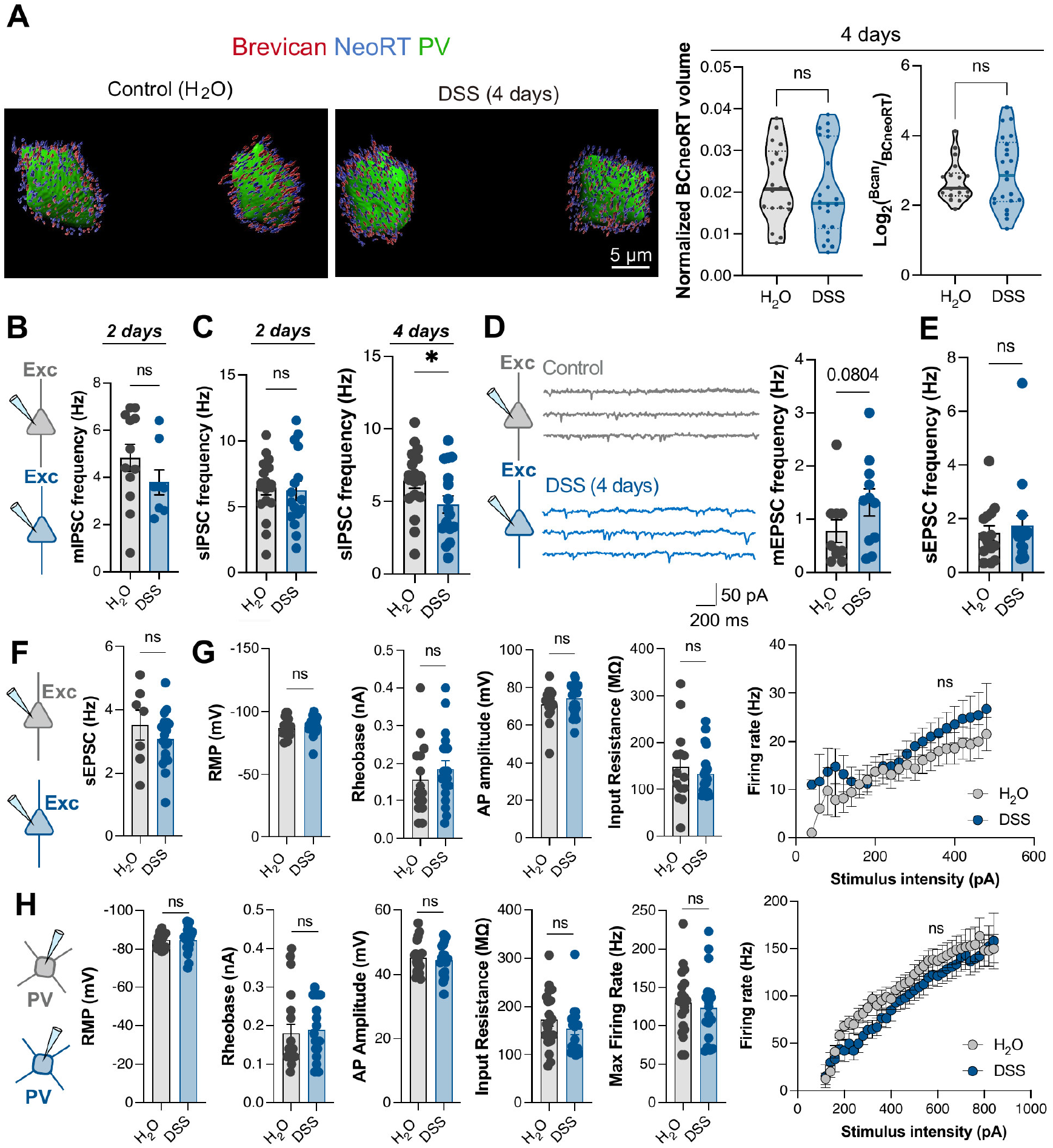
PNN proteolysis, early synaptic responses and intrinsic properties are unchanged after DSS. **(A)** Representative 3D reconstructions and quantification of ADAMTS4/5-cleaved Brevican (BCneoRT) around IC PV+ interneurons at 4 days of DSS, shown as normalized BCneoRT volume and as the log2 ratio of Brevican to BCneoRT (n = 18-22 cells from 3 mice/group). **(B)** Recording schematic and mIPSC frequency in IC excitatory neurons at 2 days of DSS (n = 8-12 cells from 3-4 mice/group). **(C)** sIPSC frequency in IC excitatory neurons at 2 and 4 days of DSS (n = 16-18 cells at 2 days and 17-18 cells at 4 days from 3-4 mice/group). **(D)** Recording schematic, representative traces and mEPSC frequency in IC excitatory neurons at 4 days of DSS (n = 10-11 cells from 3-4 mice/group). p = 0.0804. **(E)** sEPSC frequency in IC excitatory neurons at 4 days of DSS (n = 14-16 cells from 3-4 mice/group). **(F)** Recording schematic and sEPSC frequency in IC excitatory neurons in *TRAP2;Ai75* mice 5-6 days after DSS onset (n = 7-18 cells from 2 mice/group). **(G)** Intrinsic properties of IC excitatory neurons in the same mice as in F: resting membrane potential (RMP), rheobase, action potential (AP) amplitude, input resistance, and firing rate as a function of injected current; n = 15-20 cells from 2 mice/group, except for resistance where n = 14-20 cells and firing rate where n = 9-19. **(H)** Recording schematic and intrinsic properties of IC PV+ interneurons at 4 days of DSS: resting membrane potential (RMP), rheobase, action potential (AP) amplitude, input resistance, maximum firing rate, and firing rate as a function of injected current (n = 18-20 cells from 4-6 mice/group). *p<0.05; Kolmogorov-Smirnov test (A), Mann-Whitney test (B, D, E, F, RMP and input resistance in G, rheobase and input resistance in H), unpaired t-test (C, AP amplitude in G and H, rheobase in G, maximum firing rate in H), linear mixed-effects model with random intercepts and slopes per cell (firing rate in G and H). Data are mean ± SEM; in A, thick and thin lines are median and quartiles.

**Fig. S6.**
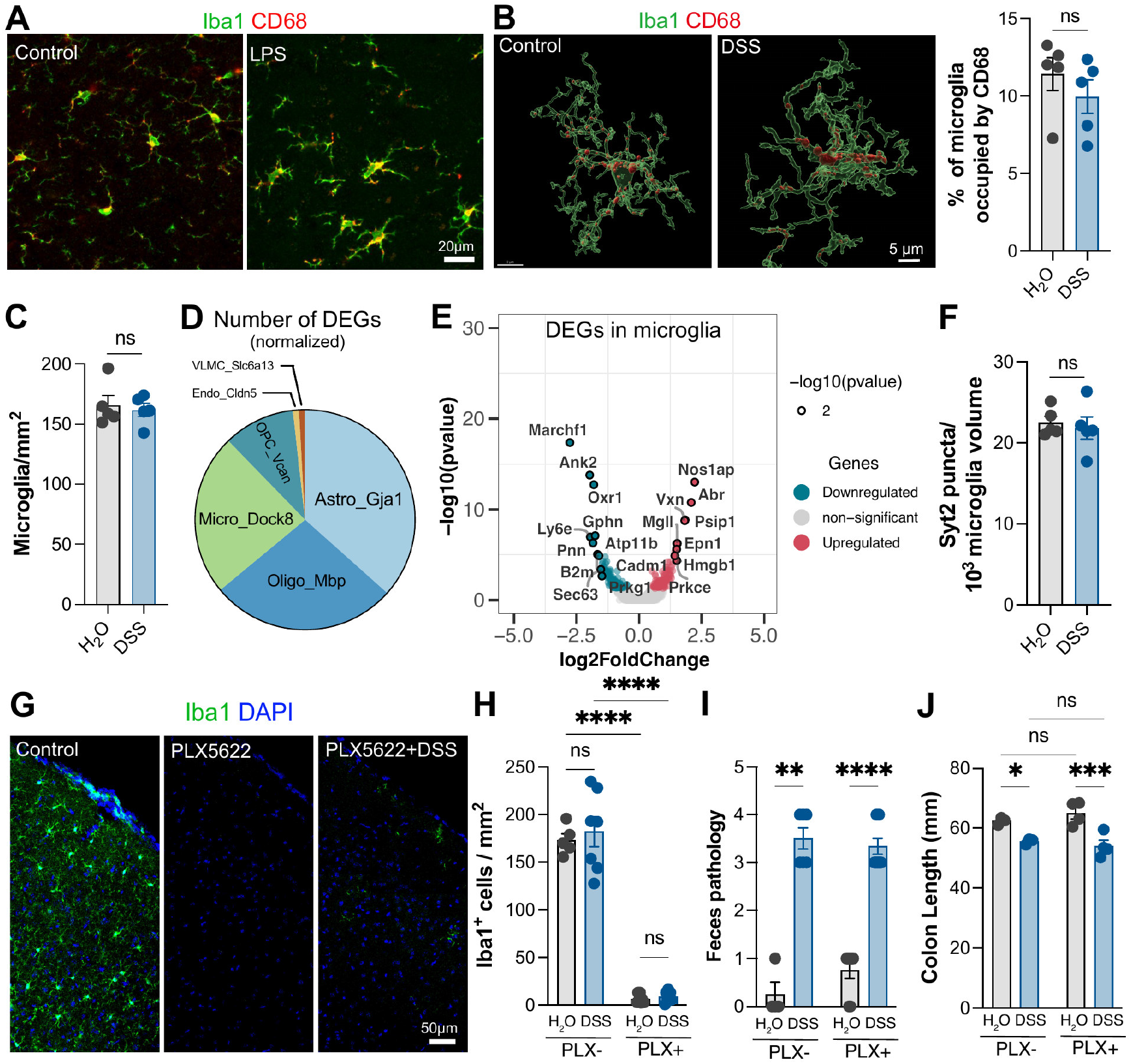
Microglia are not required for the reconfiguration of IC inhibitory circuits. **(A)** Representative images of Iba1 and CD68 staining in control and LPS-treated mice, included as a positive control for microglial activation. **(B)** Representative 3D reconstructions and fraction of microglial volume occupied by CD68+ lysosomes in controls and at 4 days of DSS (n = 5 mice/group). **(C)** Density of Iba1+ microglia in the IC of controls and at 4 days of DSS (n = 5 mice/group). **(D)** Number of differentially expressed genes (DEGs) per non-neuronal cell type, normalized to the number of nuclei recovered for that type. **(E)** Volcano plot showing DEGs in microglia at 2 days of DSS. The top 10 genes are labeled. **(F)** Density of Syt2+ puncta within microglia, normalized to microglial volume, in controls and at 4 days of DSS (n = 5 mice/group). **(G-H)** Representative images (G) and density (H) of Iba1+ microglia in the IC of control (PLX-) and PLX5622-treated (PLX+) mice on water or DSS (n = 5-8 mice/group). **(I-J)** Feces pathology score (I) and colon length (J) in PLX- and PLX+ mice on water or DSS; n = 3-4 mice/group, except in I where n = 6-9 mice/group. *p<0.05, **p<0.01, ***p<0.001, ****p<0.0001; Mann-Whitney test (B, F), unpaired t-test (C), two-way ANOVA with uncorrected Fisher’s LSD for post hoc comparisons (H, J), multiple Mann-Whitney tests (I). Data are mean ± SEM.

**Fig. S7.**
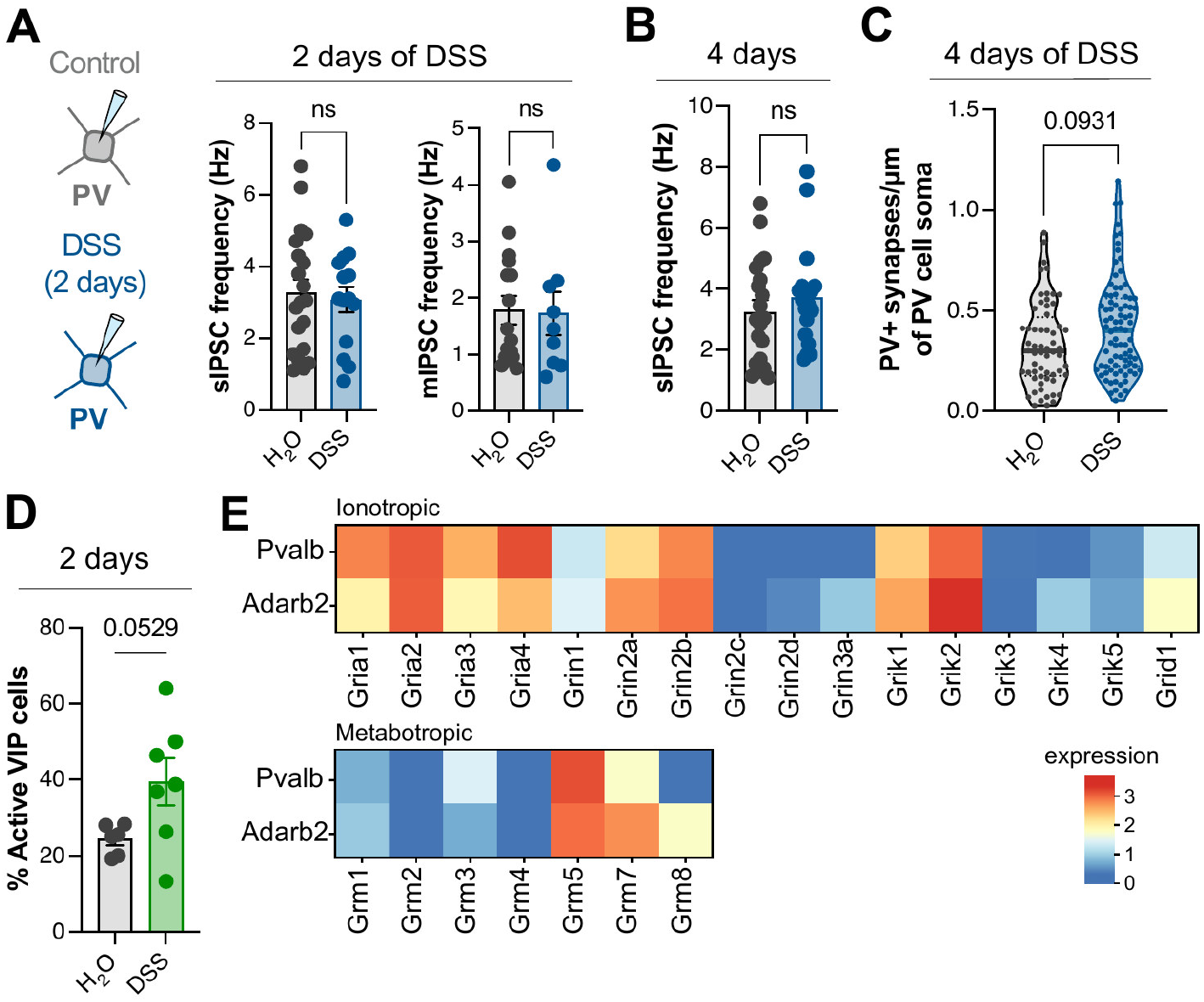
Additional inhibitory measurements onto PV+ interneurons and early VIP+ activation. **(A)** Recording schematic and sIPSC and mIPSC frequency in S5E2-GFP+ PV+ interneurons at 2 days of DSS (n = 14-20 cells for sIPSC and 9-16 cells for mIPSC from 3-5 mice/group). **(B)** sIPSC frequency in S5E2-GFP+ PV+ interneurons at 4 days of DSS; controls: n = 16-20 cells from 3 mice; DSS: n = 18-19 cells from 5 mice. **(C)** Density of PV+ (Syt2+Gephyrin+) synapses onto IC PV+ interneuron somata at 4 days of DSS (n = 60-85 cells from 3-5 mice/group). p = 0.0931. **(D)** Fraction of c-Fos+ VIP+ interneurons in the IC at 2 days of DSS (n = 5-7 mice/group). p = 0.0529. **(E)** Expression of ionotropic and metabotropic glutamate receptor genes in PV+ (Inh Pvalb) and Adarb2+ (including VIP+, Lamp5 and Sncg cells) IC interneurons (pooled across control and DSS mice). Color indicates mean normalized expression. None of these genes was differentially expressed between conditions in either population (table S2). Mann-Whitney test (A, B), Kolmogorov-Smirnov test (C), Welch’s t-test (D). Data are mean ± SEM; in C thick and thin lines are median and quartiles.

**Fig. S8.**
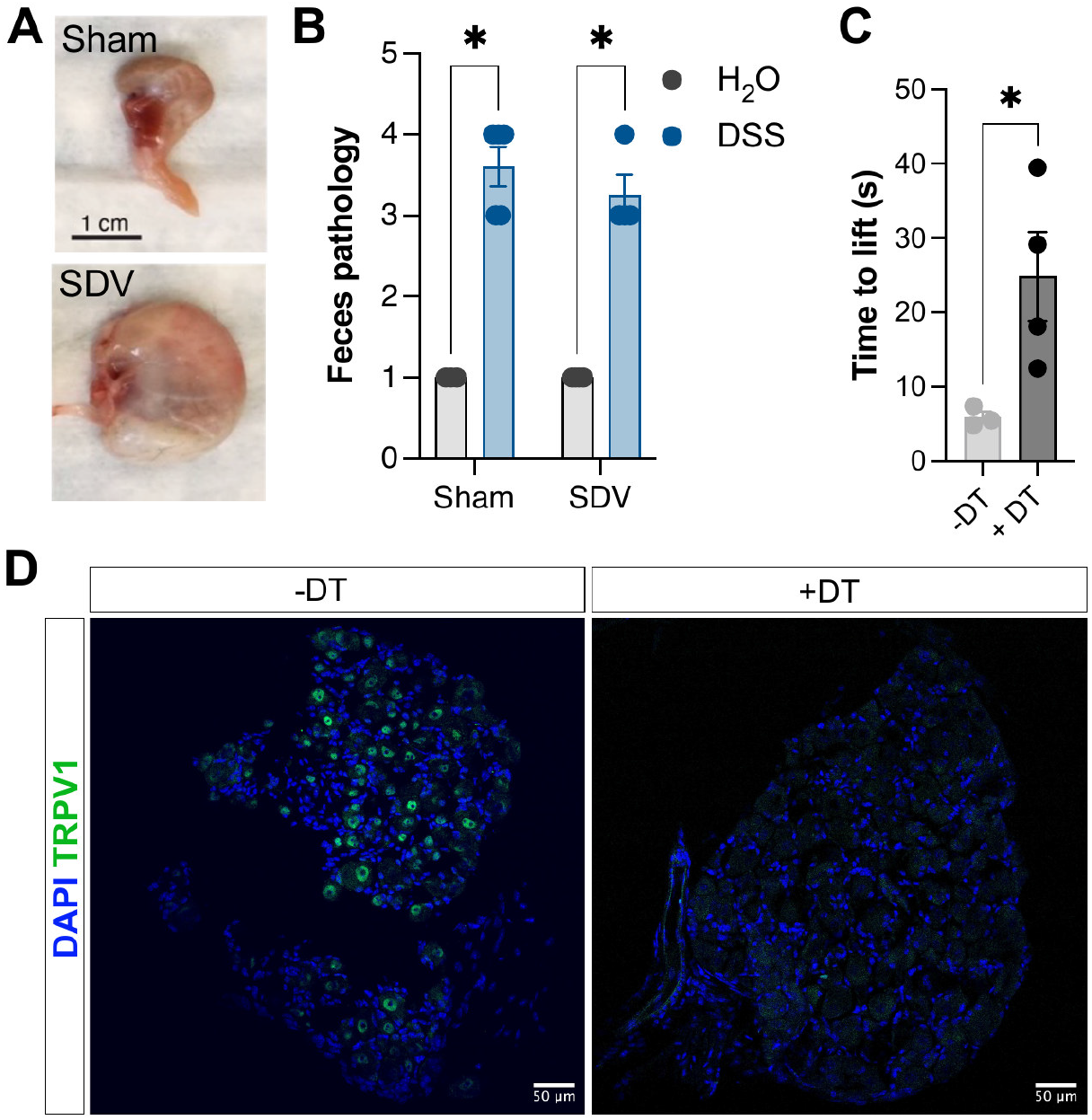
Validation of vagotomy and Trpv1+ sensory neuron ablation. (**A-B**) Representative images of the stomach confirming vagotomy (A) and feces pathology score (B) in sham and SDV mice on water or DSS (n = 3-4 mice/group). **(C-D)** Hot plate latency to hindpaw lift (C) and representative TRPV1 immunostaining in the dorsal root ganglia (D) in Trpv1-DTR mice treated with vehicle (-DT) or diphtheria toxin (+DT); n = 3-4 mice/group. *p<0.05; multiple Mann-Whitney tests (B) and Welch’s t-test (C). Data are mean ± SEM.

**Fig. S9.**
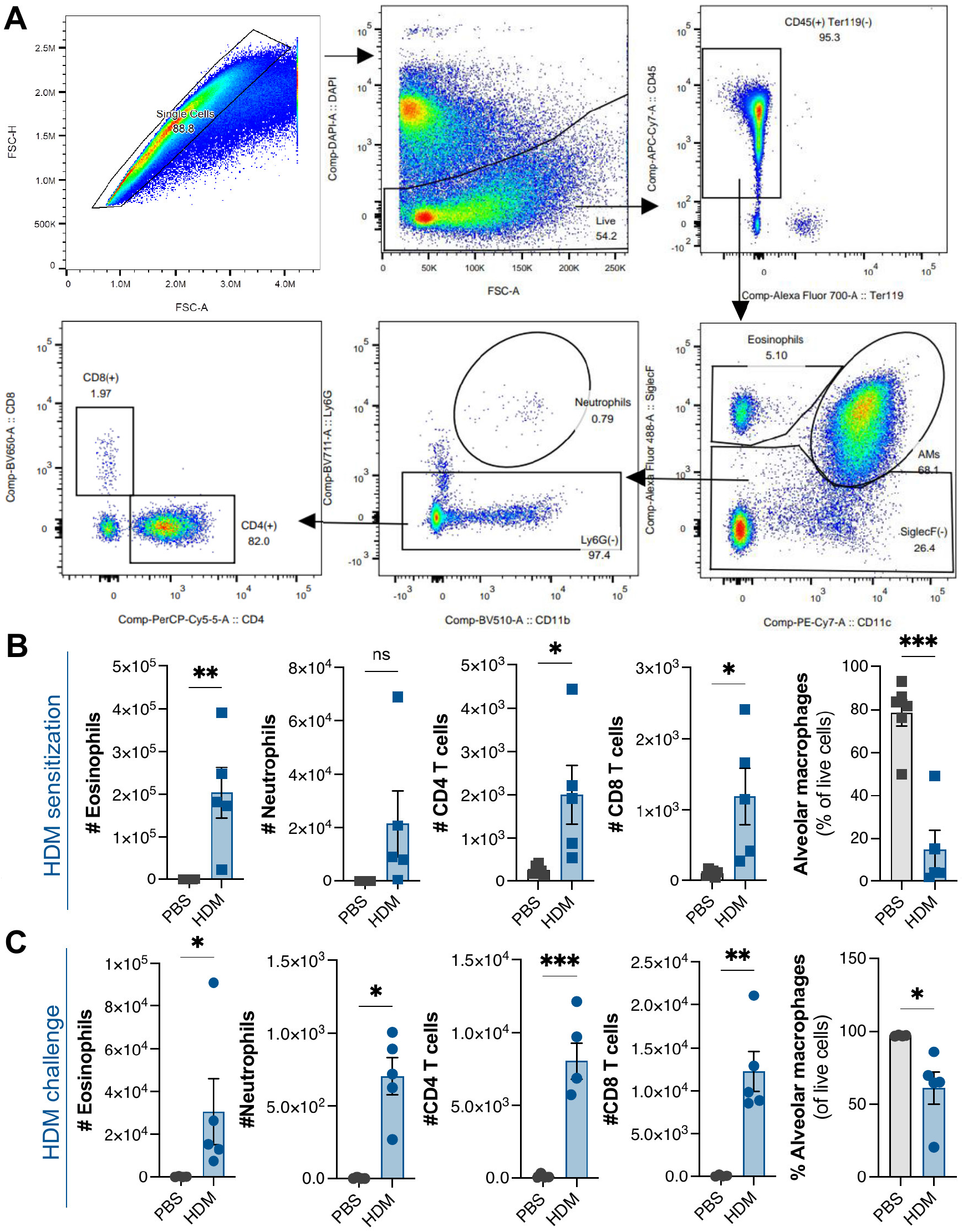
HDM sensitization and challenge induce airway inflammation. **(A)** Gating strategy for flow cytometry of bronchoalveolar lavage: single cells, live (DAPI-negative) cells, CD45+Ter119-leukocytes, SiglecF+CD11c+ alveolar macrophages and SiglecF+ eosinophils, Ly6G+CD11b+ neutrophils, and CD4+ and CD8+ T cells. **(B)** Numbers of eosinophils, neutrophils, CD4+ T cells and CD8+ T cells, and percentage of alveolar macrophages in bronchoalveolar lavage after HDM sensitization (n = 5-6 mice/group). **(C)** As in (B), after HDM challenge (n = 3-4 mice/group). *p<0.05, **p<0.01, ***p<0.001; unpaired t-test, except for neutrophils and alveolar macrophages in B, where Welch’s t-test was used. Data are mean ± SEM.

**Fig. S10.**
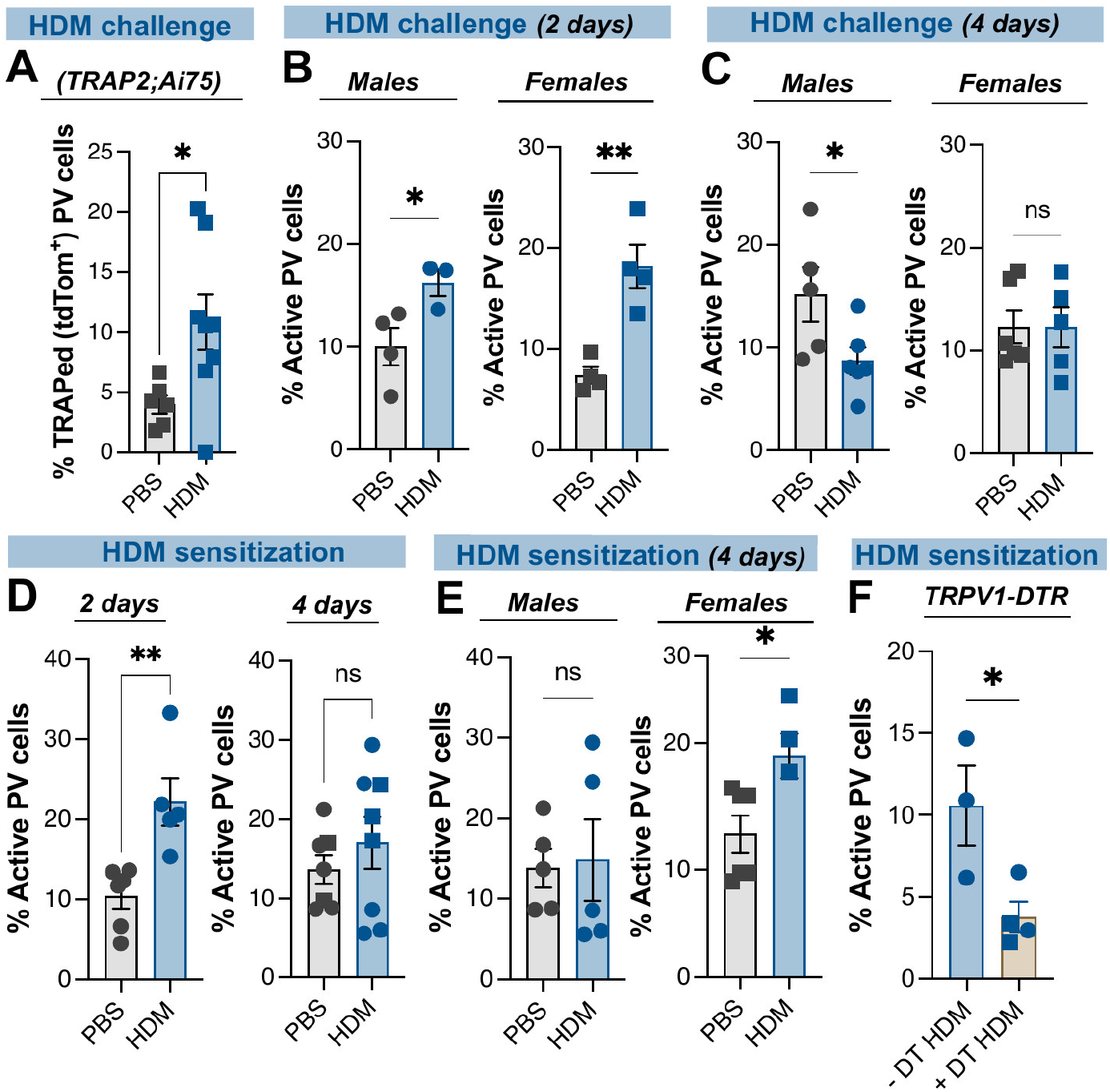
PV+ interneuron responses to HDM across sexes and paradigms. **(A)** Fraction of TRAPed (tdTomato+) PV+ interneurons in the IC of *TRAP2;Ai75* female mice after HDM challenge, with tamoxifen given at the time of challenge and brain collection at 3 days post-challenge (n = 6-8 mice/group). **(B)** Fraction of c-Fos+ PV+ interneurons in the IC of male and female mice 2 days after HDM challenge (n = 3-4 mice/group). Data are the same as in Fig. 4J but split by sex. **(C)** Fraction of c-Fos+ PV+ interneurons in the IC of male and female mice 4 days after HDM challenge (n = 5-6 mice/group). Male data are the same as in Fig. 4J but shown next to females for comparison. **(D)** Fraction of c-Fos+ PV+ interneurons in the IC 2 and 4 days after HDM sensitization (n = 5-6 mice/group for the 2-day time point and 7-8 mice/group for the 4-day time point). **(E)** Fraction of c-Fos+ PV+ interneurons in the IC of male and female mice 4 days after HDM sensitization (n = 3-5 mice/group). Data are the same as in D but split by sex. **(F)** Fraction of c-Fos+ PV+ interneurons in the IC 2 days after HDM sensitization in *Trpv1-DTR* mice treated with vehicle (-DT) or diphtheria toxin (+DT); n = 3-4 mice/group. *p<0.05, **p<0.01; unpaired t-test. Data are mean ± SEM; in this figure, circles and square symbols indicate males and females, respectively.

**Fig. S11.**
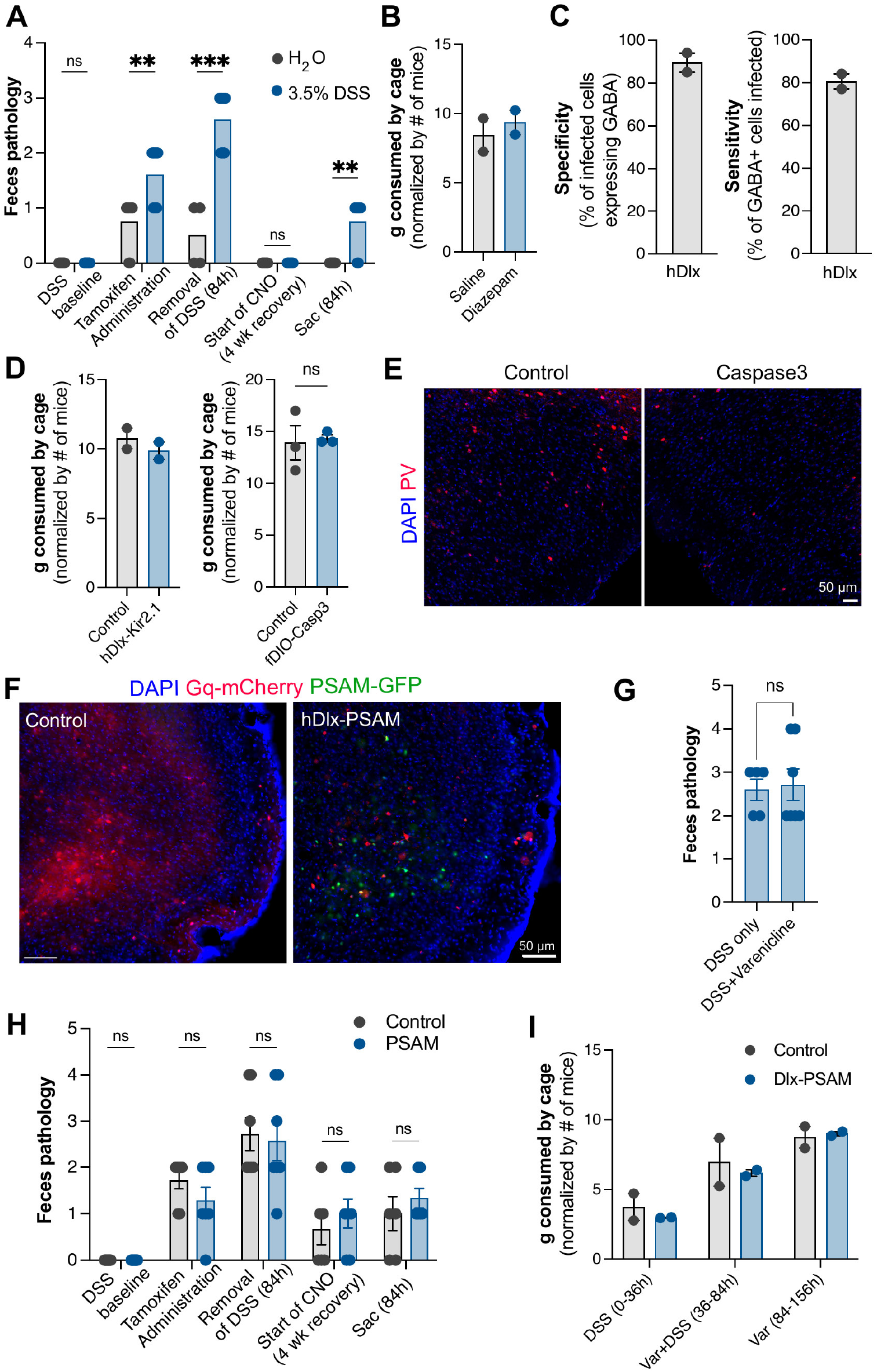
Controls for the immune engram manipulation experiments. **(A)** Feces pathology score across the experimental timeline in control and 3.5% DSS mice (n = 5 mice/group). **(B)** Drinking solution consumed per cage, normalized to the number of mice, in saline- and diazepam-treated mice (n = 2-3 cages/group). **(C)** Specificity (percentage of infected cells expressing GABA) and sensitivity (percentage of GABA+ cells infected) of the hDlx enhancer in the IC (n = 2 mice). **(D)** Drinking solution consumed per cage, normalized to the number of mice, in control and hDlx-Kir2.1 mice and in control and fDIO-Caspase3 mice (n = 2-3 cages/group). **(E)** Representative PV immunostaining in the IC of control and fDIO-Caspase3 mice. **(F)** Representative images of Gq-mCherry and hDlx-PSAM-GFP expression in the IC of control and hDlx-PSAM mice. **(G)** Feces pathology score across the experimental timeline in control and hDlx-PSAM mice (n = 5-7 mice/group). **(H)** Feces pathology score in mice on DSS alone or DSS with varenicline (n = 5-7 mice/group). **(I)** Drinking solution consumed per cage, normalized to the number of mice, in control and hDlx-PSAM mice (n = 2-3 cages/group). **p<0.01, ***p<0.001; multiple Mann-Whitney tests (A, G) and Mann-Whitney test (B, D, H, I). Data are mean ± SEM.

## Notes

### Competing Interest Statement

The authors have declared no competing interest.

